# A Versatile Linker for Probes Targeting Hydrolases via *In Situ* labeling

**DOI:** 10.1101/2021.06.14.448363

**Authors:** Jun Liu, Zixin Chen, Chao Cui, Ashton L. Sigler, Lina Cui

## Abstract

Hydrolases are important molecules that are involved in a wide range of biological functions and their activities are tightly regulated in healthy or diseased states. Detecting or imaging the activities of hydrolases, therefore, can reveal underlying molecular mechanisms in the context of cells to organisms, and their correlation with different physiological conditions can therefore be used in diagnosis. Due to the nature of hydrolases, substrate-based probes can be activated in their catalytic cycles, and cleavage of covalent bonds frees reporter moieties. For test-tube type bulk detection, spatial resolution is not a measure of importance, but for cell- or organism-based detection or imaging, spatial resolution is a key factor for probe sensitivity that influences signal-to-background ratio. One strategy to improve spatial resolution of the probes is to form a covalent linkage between the reporter moiety and intracellular proteins upon probe activation by the enzyme. In this work, we developed a generalizable linker chemistry that would allow *in situ* labeling of various imaging moieties via quinone methide species. To do so, we synthesized probes containing a monofluoromethyl or a difluoromethyl groups for β-galactosidase activation, while using fluorescein as a fluorescent reporter. The labeling efficacy of these two probes was evaluated *in vitro*. The probe bearing a monofluormethyl group exhibited superior labeling efficiency in imaging β-galactosidase activity in living cells. This study provides a versatile linker for applying quinone methide chemistry in the development of hydrolase-targeting probes involving *in situ* labeling.

## Introduction

Molecular imaging probes have become important tools for interrogating biological functions both spatially and temporally in live cells or *in vivo*.^1^ Hydrolases, enzymes that catalyze the hydrolysis of a covalent bond, are important molecules that are involved in a wide range of biological functions, and, like other biomolecules, their activities are tightly regulated in healthy or diseased states. Detecting or imaging the activities of hydrolases, therefore, can reveal underlying molecular mechanisms in the context of cells to organisms, and their correlation with different physiological conditions can therefore be used in diagnosis.^2^ Due to the nature of hydrolases, substrate-based probes can be activated in their catalytic cycles, and cleavage of covalent bonds frees reporter moieties that provide either positive or negative signal output, and they have been used widely to assess enzyme activities in cells and *in vivo* due to sufficient cell uptake, high specificity, and tissue distribution.^3^ A major limitation of most of these small molecule imaging probes is that they can easily diffuse away from the site of activation, resulting in poor sensitivity and low signal-to-background ratio.^5^ For test-tube type bulk detection, spatial resolution is not a measure of importance, but for cell- or organism-based detection or imaging, spatial resolution is a key factor for probe sensitivity that influences signal-to-background ratio. Various strategies have been developed to improve the spatial resolution.^6-8^ For example, small molecule probes can be activated by the target enzyme, and due to the change in hydrophobicity before and after enzyme activation, they can self-assembly into nanometersized particles or aggregates and retain at site of activation.^9-16^ Another strategy is to form a covalent bond between the reporter moiety and intracellular proteins via the reaction between a labile electrophilic moiety on the imaging probe and nucleophiles such as thiols or amines on proteins.^6, 17-20^

Quinone methide (QM) chemistry has been explored to form covalent bonds between the fluorescent probe and surrounding proteins.^21-24^ Quinone methide is an unstable intermediate that is susceptible to nucleophilic attack.^25, 26^ It can be generated by external stimuli under physiological conditions from a stable masked phenol precursor bearing a leaving group (such as a fluorine atom) on the methyl group at the para- or ortho-position.^27^ This strategy has been used in inhibitors and prodrugs, ^28-30^ self-immolative linkers in dendrimers,^31^ chemical selection of catalytic antibodies, ^32,33^ activity-based probes for various enzymes, including phosphatase,^34^ glycosidases,^22, 23, 35-41^, tyrosine phosphatase,^42^ steroid sulfatase,^43^ and beta-lactamase,^44^ cell imaging^45, 46^ and probing protein-protein/DNA interactions.^47^ Among these applications, the fluorine on either the monofluoromethyl or difluoromethyl group was used as a leaving group to form the active o-quinone methide intermediate via spontaneous elimination of a molecule of HF. However, the application of the quinone methide chemistry has been dwarfed by the dilemma that the monofluoromethyl group is more reactive and the protein-conjugated products are more stable than its difluoromethyl counterpart, but molecules containing a monofluoromethyl group suffer from poor stability in aqueous media, as the reactive monofluoromethyl group tends to be hydrolyzed by the surrounding water before the probe molecule reaches the active site of the target protein. Some were able to obtain stable molecules containing a monofluoromethyl group ^45,48^, while many others, including us, failed.^49, 50^ Herein, we seek to develop a linker chemistry can provide a generalizable approach for the use of quinone methide in probe development. We designed and synthesized three molecules, one with a *N-* (4-hydroxyphenyl)imidothiocarbamate linker, and the other two incorporated with either a monofluoromethyl or a difluoromethyl group on the phenol group (Figure 1). All probe molecules contained a fluorescein (FITC) at its “on” state as the reporter. Without fluorescence change after target activation with these FITC probes, our approach provided direct visual comparison of the labeling efficiency of these molecules both *in vitro* and in live cells.

**Figure 1.**
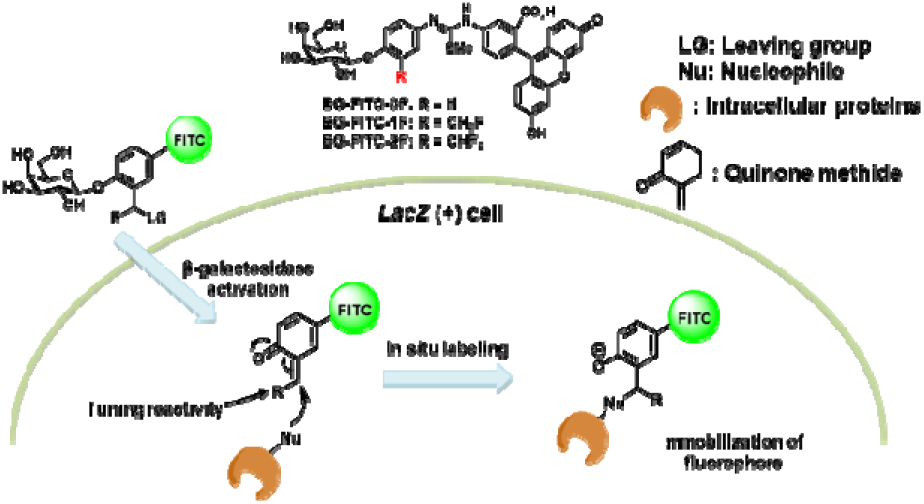
Schematic illustration of β-galactosidase-sensitive fluorescent probe immobilized by intracellular proteins via quinone methide

## Materials and Methods

### Chemical Synthesis

All chemicals were purchased from commercial sources unless otherwise noted. High-resolution mass spectrometry (HRMS) samples were analyzed on a Waters LCT Premier Mass Spectrometer. High-performance liquid chromatography (HPLC) was performed on a Dionex Ultimate 300 HPLC System (Thermo Scientific) equipped with a GP50 gradient pump and an in-line diode array UV-Vis detector. Reverse-phase C18 columns were used with acetonitrile/water gradient mobile phase containing 0.1% trifluoroacetic acid. NMR spectra were recorded on Bruker instruments (500 MHz and 600 MHz for ^1^H NMR, 126 MHz and 151 MHz for ^13^C NMR) and internally referenced to the residual solvent signals (^1^H: δ 7.26; ^13^C: δ 77.16 for CDCl_3_, ^1^H: δ 3.31; ^13^C: δ 49.0 for CD_3_OD respectively). NMR chemical shifts (δ) and the coupling constants (J) for ^1^H and ^13^C NMR are reported in parts per million (ppm) and in Hertz, respectively. The following conventions are used for multiplicities: s, singlet; d, doublet; t, triplet; m, multiplet; and dd, doublet of doublet.

### SDS-PAGE Analysis of FITC-Labeled β-galactosidase

10 μg of β-galactosidase from *Escherichia coli* (Millipore Sigma) in 20 μL 1X PBS buffer was incubated with different concentrations (1, 5 or 10 μM) of **BG-FITC-0F**, **BG-FITC-1F** or **BG-FITC-2F** for 1 hour at 37 °C. The samples were then subjected to sodium dodecyl sulphate-polyacrylamide gel electrophoresis (SDS-PAGE, Bio-Rad Mini-PROTEAN^®^ TGX™ 4-15% Gel) according to manufacturer’s protocols for electrophoresis at 110 V for 70 min. Both fluorescent image and Coomassie staining of the gel were taken using Bio-Rad Gel-Doc. The fluorescence intensity of FITC-labeled β-galactosidase was quantified by ImageJ.

### Flow Cytometry

CT26.WT and CT26.CL25 (*LacZ+*) cells were purchased from ATCC. CT26.WT and CT26.CL25 cells were seeded in 24-well plates at a density of 2× 10^4^ cells/well until they reach 60-70% confluency. Then the cells were incubated with 10 μM of **BG-FITC-0F**, **BG-FITC-1F** or **BG-FITC-2F** for 4 or 14 hours at 37 °C. The cells were then washed three times with cold PBS buffer, digested with 0.25% trypsin, and collected by centrifugation at 100×g for 5 min. The resulting cell pellets were resuspended and fixed with 4% paraformaldehyde in PBS, followed by washing with PBS buffer and analyzed using flow cytometer (Accuri C6, Becton Dickinson, USA).

### Fluorescence Imaging of Cells

CT26.WT and CT26.CL25 cells were seeded onto poly-L-lysine-coated cover glasses in 24-well plates at a density of 2×10^4^ cells/well until they reached 60-70% confluency. Then the cells were incubated with 10 μM of **BG-FITC-0F**, **BG-FITC-1F** or **BG-FITC-2F** for 14 hours at 37 °C. The cells were then washed three times with cold PBS buffer followed by fixation using 4% paraformaldehyde. Lastly, the cells were stained with DAPI for 5 min, washed with PBS and were mounted onto tissue slides by ProLong Gold Antifade Mountant (Invitrogen, USA). The fluorescence images were acquired using a Nikon Eclipse Ti2 fluorescence microscope with the DAPI and FITC filters.

## Results and Discussions

### Probe Design and Synthesis

We designed three probes each with a β-galactoside moiety linked to a fluorescein (FITC) fluorescent reporter via a phenol linker (Figure 1). **BG-FITC-0F** has a *N*-(4-hydroxyphenyl)imidothiocarbamate linker, and **BG-FITC-1F** and **BG-FITC-2F** carry a monofluoromethyl or difluoromethyl groups at the ortho position on the phenol. Having fluoro-substitution at this position allows the formation of a reactive quinone methide intermediate once the β-galactoside is cleaved by β-galactosidase. Monofluoromethyl and difluoromethyl groups have been shown to be good trapping groups with other activity-based probes ^24, 28, 45^. The quinone methide intermediate can then react with nucleophiles such as amines or thiols that are ubiquitous on intracellular proteins and form covalent linker between FITC and intracellular proteins. The isothiourea linker provides increased stability ^23^. **BG-FITC-0F** does not have such trapping unit thus would not form covalent linker with intracellular proteins and serves as a control probe in this study. These probes are constantly on fluorescent probes thus can provide accurate information on labeling efficiency unaffected by change in fluorescence. The synthesis of these three probes are described in detail in the supporting information. It is important to note that all three probes, including the monofluoromethyl group containing **BG-FITC-1F**, are very stable in aqueous media, and they all have a long (tested for several months) shelf-life.

### Probe Labeling Efficiency and Gel Analysis

With the three probes in hand, we next evaluated their labeling efficiency with recombinant β-galactosidase. We incubated 10 μg of β-galactosidase with 1, 5 or 10 μM of each of the three probes. The resulting mixtures were analyzed by SDS-PAGE gel and the fluorescence intensities of FITC-labeled β-galactosidase were compared parallelly. As shown in Figure 2, compared to the control probe **BG-FITC-0F**, both **BG-FITC-1F** and **BG-FITC-2F** showed clear labeling of β-galactosidase at 10 μM. This result is consistent with the theory that both monofluoromethyl and difluoromethyl groups at the ortho position on the phenol linker can act as trapping units that immobilize the fluorophore onto proteins. While comparing the labeling efficiency at lower concentrations, **BG-FITC-1F** showed more efficient labeling than **BG-FITC-2F**. The different labeling efficiency between these two structural analogues can be attributed to the different reactivity of monofluoromethyl and difluoromethyl groups ^35, 44^. After enzymatic cleavage, the active quinone methide intermediate is conjugated to proteins with a stable covalent bond for **BG-FITC-1F**, whereas the adduct of protein with substrate **BG-FITC-2F** is less stable due to the existence of the second C-F bond. This unstable protein-probe adduct can generate quinone methide intermediate again via the elimination of another molecule of HF, followed by nucleophilic attack from the surrounding water molecules to afford an unstable intermediate that further hydrolyzes and releases the free fluorophore, lowering the labeling efficiency ^44, 51^.

**Figure 2.**
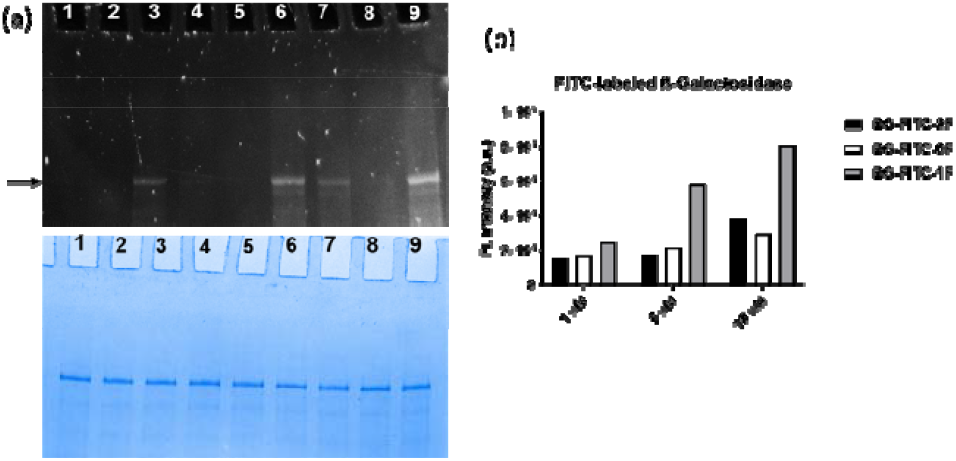
(a) SDS-PAGE analysis of FITC-labeled β-Galactosidase (10 μg) after it was incubated with different concentrations of BG-FITC-2F, BG-FITC-0F or BG-FITC-1F. Lane 1-3: 1 μM of BG-FITC-2F, BG-FITC-0F or BG-FITC-1F; Lane 4-6: 5 μM of BG-FITC-2F, BG-FITC-0F or BG-FITC-1F; Lane 7-8: 1 μM of BG-FITC-2F, BG-FITC-0F or BG-FITC-1F. (b) Quantification of fluorescence intensity of FITC-labeled β-galactosidase (indicated by arrow in figure (a)) in each lane.

### Flow Cytometry and Cell Imaging

We further characterized the labeling efficiency of these probes in mouse colon cancer cell line CT26. CT26.CL25 is stably transduced with the *lacZ* gene to express β-galactosidase and the wildtype CT26.WT was used as control. The cells were first incubated with 10 μM of each of the three probes for 4 or 14 hours and the resulting cells were analyzed by flow cytometry. As shown in Figure 3, both **BG-FITC-1F** and **BG-FITC-2F** showed significant labeling in CT26.CL25 compared to the control probe **BG-FITC-0F**. **BG-FITC-1F** also showed significantly higher labeling efficiency compared to **BG-FITC-2F**. These results were consistent with what we observed in the SDS-PAGE gel analysis. We also performed fluorescence cell imaging of CT26.WT and CT26.CL25 after incubating with the probes for 14 hours. As shown in Figure 4, cell imaging also demonstrated a similar result as the flow cytometry with **BG-FITC-1F** showing the highest labeling efficiency.

**Figure 3.**
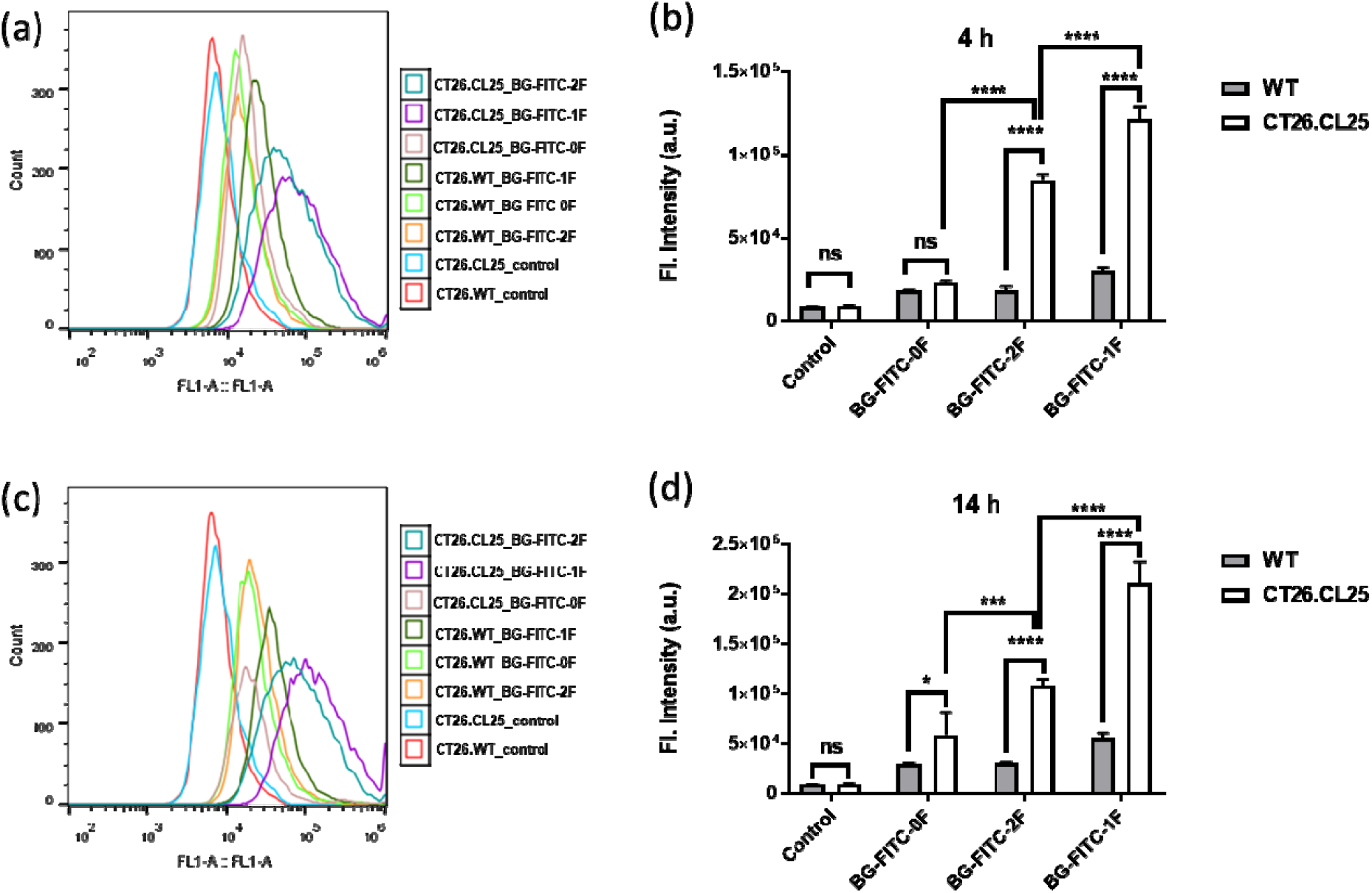
(a) FACS analysis of CT26.WT and CT26.CL25 cells after incubation with 10 μM of BG-FITC-0F, BG-FITC-1F or BG-FITC-2F for 4 hours. (b) Quantification of average fluorescence in (a). (c) FACS analysis of CT26.WT and CT26.CL25 cells after incubation with 10 μM of BG-FITC-0F, BG-FITC-1F or BG-FITC-2F for 14 hours. (d) Quantification of average fluorescence in (c). NS, not significant; *, P<0.1; ***, P<0.001; ****, P<0.0001.

**Figure 4.**
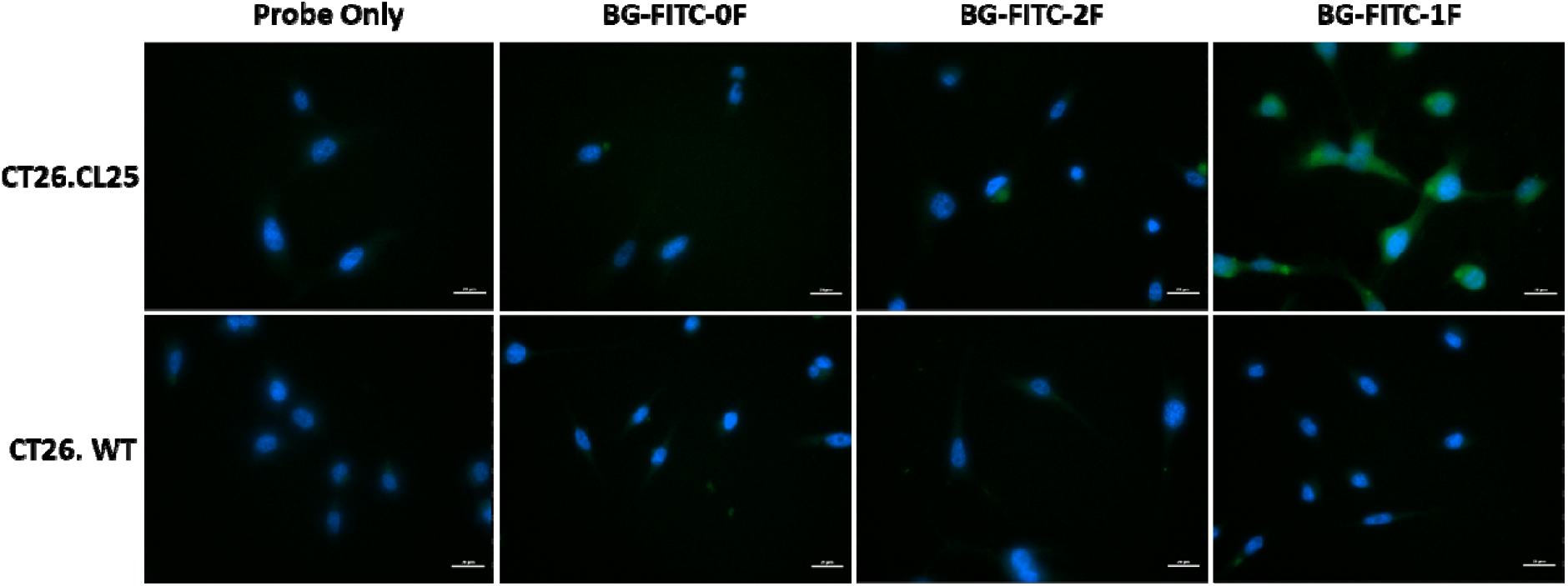
Fluorescent cell images of CT26.WT and CT26.CL25 cells after incubation with 10 μM of BG-FITC-0F, BG-FITC-1F or BG-FITC-2F for 14 hours. Scale bar: 20 μm.

## Conclusions

While quinone methide chemistry is very desirable in the field of chemical biology, drug discovery and molecular imaging, its application has been impeded by the difficulty in choosing proper quinone methide species. Monofluoromethyl group is very appealing in many cases for its rapid labeling reaction with the target protein and the formation of stable protein-probe conjugates, but probes containing a monofluoromethyl group tend to be hydrolyzed too fast to be used in any biological applications. In this work, we developed a generalizable linker chemistry that would allow *in situ* labeling of various imaging moieties via quinone methide species. For this purpose, we designed and synthesized three fluorescent imaging probes, each carrying a β-galactosidase-sensitive caging unit, a phenol linker with or without a in aqueous media, rendering their suitability in biological applications. The labeling efficiency of these probes was compared through SDS-PAGE gel analysis, flow cytometry and fluorescence cell imaging. The results demonstrated that the monofluorinated substrate has a superior labeling efficiency due to the formation of a more stable covalent bond between the fluorescent reporter and surrounding proteins, producing improved long-term signal retention and imaging sensitivity. This linker we explored in this work can be used in various other applications where FITC is replaced by functional moieties of choice.

## Data Availability Statement

Details for chemical synthesis and compound characterizations are included in the supporting material.

## Author Contributions

J.L. and L.C. conceived and designed the project. J.L. performed all the chemical synthesis and compound characterization. Z.C., C.C., and A.L.S. performed gel electrophoresis, fluorescence microscopy, and flow cytometry. J.L., Z.C., and L.C. wrote the paper, and all authors commented on the paper.

## References

1. Weissleder, R.; Mahmood, U., Molecular imaging. Radiology 2001, 219 (2), 316–333.

2. Yang, C.; Wang, Q.; Ding, W., Recent progress in the imaging detection of enzyme activities in vivo. RSC Advances 2019, 9 (44), 25285–25302.

3. Li, S.-D.; Huang, L., Pharmacokinetics and biodistribution of nanoparticles. Molecular Pharmaceutics 2008, 5 (4), 496–504.

4. Owensiii, D.; Peppas, N., Opsonization, biodistribution, and pharmacokinetics of polymeric nanoparticles. International Journal of Pharmaceutics 2006, 307 (1), 93–102.

5. Johnson, I., Review: Fluorescent probes for living cells. The Histochemical Journal 1998, 30 (3), 123–140.

6. Edgington, L. E.; Verdoes, M.; Bogyo, M., Functional imaging of proteases: recent advances in the design and application of substrate-based and activity-based probes. Current Opinion in Chemical Biology 2011, 15 (6), 798–805.

7. Qi, G.-B.; Gao, Y.-J.; Wang, L.; Wang, H., Self-assembled peptide-based nanomaterials for biomedical imaging and therapy. Advanced Materials 2018, 30 (22).8.

8. Hai, Z.; Liang, G., Intracellular self-assembly of nanoprobes for molecular imaging. Advanced Biosystems 2018, 2 (8).

9. Gao, Y.; Shi, J.; Yuan, D.; Xu, B., Imaging enzyme-triggered self-assembly of small molecules inside live cells. Nature Communications 2012, 3 (1).

10. Liang, G.; Ren, H.; Rao, J., A biocompatible condensation reaction for controlled assembly of nanostructures in living cells. Nature Chemistry 2009, 2 (1), 54–60.

11. Ye, D.; Shuhendler, A. J.; Cui, L.; Tong, L.; Tee, S. S.; Tikhomirov, G.; Felsher, D. W.; Rao, J., Bioorthogonal cyclization-mediated in situ self-assembly of small-molecule probes for imaging caspase activity in vivo. Nature Chemistry 2014, 6 (6), 519–526.

12. Onogi, S.; Shigemitsu, H.; Yoshii, T.; Tanida, T.; Ikeda, M.; Kubota, R.; Hamachi, I., In situ real-time imaging of self-sorted supramolecular nanofibres. Nature Chemistry 2016, 8 (8), 743–752.

13. Li, J.; Gao, Y.; Kuang, Y.; Shi, J.; Du, X.; Zhou, J.; Wang, H.; Yang, Z.; Xu, B., Dephosphorylation of d-peptide derivatives to form biofunctional, supramolecular nanofibers/hydrogels and their potential applications for intracellular imaging and intratumoral chemotherapy. Journal of the American Chemical Society 2013, 135 (26), 9907–9914.

14. Chen, Z.; Chen, M.; Cheng, Y.; Kowada, T.; Xie, J.; Zheng, X.; Rao, J., Exploring the condensation reaction between aromatic nitriles and amino thiols to optimize in situ nanoparticle formation for the imaging of proteases and glycosidases in cells. Angewandte Chemie 2020, 132 (8), 3298–3305.

15. Chen, Z.; Chen, M.; Zhou, K.; Rao, J., Pre targeted imaging of protease activity through in situ assembly of nanoparticles. Angewandte Chemie International Edition 2020, 59 (20), 7864–7870.

16. Schleyer, K. A.; Datko, B. D.; Burnside, B.; Cui, C.; Ma, X.; Grey, J. K.; Cui, L., Responsive fluorophore aggregation provides spectral contrast for fluorescence lifetime imaging. ChemBioChem 2020, 21 (15), 2196–2204.

17. Blum, G.; von Degenfeld, G.; Merchant, M. J.; Blau, H. M.; Bogyo, M., Noninvasive optical imaging of cysteine protease activity using fluorescently quenched activity-based probes. Nature Chemical Biology 2007, 3 (10), 668–677.

18. Blum, G.; Mullins, S. R.; Keren, K.; Fonovič, M.; Jedeszko, C.; Rice, M. J.; Sloane, B. F.; Bogyo, M., Dynamic imaging of protease activity with fluorescently quenched activity-based probes. Nature Chemical Biology 2005, 1 (4), 203–209.

19. Edgington, L. E.; Berger, A. B.; Blum, G.; Albrow, V. E.; Paulick, M. G.; Lineberry, N.; Bogyo, M., Noninvasive optical imaging of apoptosis by caspase-targeted activity-based probes. Nature Medicine 2009, 15 (8), 967–973.

20. Kwan, D. H.; Chen, H. M.; Ratananikom, K.; Hancock, S. M.; Watanabe, Y.; Kongsaeree, P. T.; Samuels, A. L.; Withers, S. G., Self-immobilizing fluorogenic imaging agents of enzyme activity. Angew. Chem. Int. Ed. 2011, 50 (1), 300–3.

21. Turner, A. B., Quinone methides. Quarterly Reviews, Chemical Society 1964, 18 (4).

22. Kurogochi, M.; Nishimura, S.; Lee, Y. C., Mechanism-based fluorescent labeling of beta-galactosidases. An efficient method in proteomics for glycoside hydrolases. The Journal of biological chemistry 2004, 279 (43), 44704–12.

23. Cheng, T. C.; Roffler, S. R.; Tzou, S. C.; Chuang, K. H.; Su, Y. C.; Chuang, C. H.; Kao, C. H.; Chen, C. S.; Harn, I. H.; Liu, K. Y.; Cheng, T. L.; Leu, Y. L., An activity-based near-infrared glucuronide trapping probe for imaging beta-glucuronidase expression in deep tissues. J Am Chem Soc 2012, 134 (6), 3103–10.

24. Song, H.; Li, Y.; Chen, Y.; Xue, C.; Xie, H., Highly Efficient Multiple-Labeling Probes for the Visualization of Enzyme Activities. Chemistry – A European Journal 2019, 25 (61), 13994–14002.

25. Bai, W. J.; David, J. G.; Feng, Z. G.; Weaver, M. G.; Wu, K. L.; Pettus, T. R., The domestication of ortho-quinone methides. Acc Chem Res 2014, 47 (12), 3655–64.

26. Singh, M. S.; Nagaraju, A.; Anand, N.; Chowdhury, S., ortho-Quinone methide (o-QM): a highly reactive, ephemeral and versatile intermediate in organic synthesis. RSC Adv. 2014, 4 (99), 55924–55959.

27. Minard, A.; Liano, D.; Wang, X.; Di Antonio, M., The unexplored potential of quinone methides in chemical biology. Bioorg Med Chem 2019, 27 (12), 2298–2305.

28. Chiba, M.; Kamiya, M.; Tsuda-Sakurai, K.; Fujisawa, Y.; Kosakamoto, H.; Kojima, R.; Miura, M.; Urano, Y., Activatable photosensitizer for targeted ablation of lacZ-positive cells with singlecell resolution. ACS Cent. Sci. 2019, 5 (10), 1676–1681.

29. Myers, J.; Widlanski, T., Mechanism-based inactivation of prostatic acid phosphatase. Science 1993, 262 (5138), 1451–1453.

30. Haba, K.; Popkov, M.; Shamis, M.; Lerner, R. A.; Barbas, C. F.; Shabat, D., Single-triggered trimeric prodrugs. Angewandte Chemie International Edition 2005, 44 (5), 716–720.

31. Gnaim, S.; Shabat, D., Quinone-methide species, a gateway to functional molecular systems: from self-immolative dendrimers to long-wavelength fluorescent dyes. Accounts of Chemical Research 2014, 47 (10), 2970–2984.

32. Janda, K. D.; Lo, L. C.; Lo, C. H.; Sim, M. M.; Wang, R.; Wong, C. H.; Lerner, R. A., Chemical selection for catalysis in combinatorial antibody libraries. Science 1997, 275 (5302), 945–8.

33. Cesaro-Tadic, S.; Lagos, D.; Honegger, A.; Rickard, J. H.; Partridge, L. J.; Blackburn, G. M.; Pluckthun, A., Turnover-based in vitro selection and evolution of biocatalysts from a fully synthetic antibody library. Nature biotechnology 2003, 21 (6), 679–85.

34. Betley, J. R.; Cesaro-Tadic, S.; Mekhalfia, A.; Rickard, J. H.; Denham, H.; Partridge, L. J.; Pluckthun, A.; Blackburn, G. M., Direct screening for phosphatase activity by turnover-based capture of protein catalysts. Angew Chem Int Ed Engl 2002, 41 (5), 775–7.

35. Hsu, Y. L.; Nandakumar, M.; Lai, H. Y.; Chou, T. C.; Chu, C. Y.; Lin, C. H.; Lo, L. C., Development of activity-based probes for imaging human alpha-l-fucosidases in cells. The Journal of organic chemistry 2015, 80 (16), 8458–63.

36. Chauvigne-Hines, L. M.; Anderson, L. N.; Weaver, H. M.; Brown, J. N.; Koech, P. K.; Nicora, C. D.; Hofstad, B. A.; Smith, R. D.; Wilkins, M. J.; Callister, S. J.; Wright, A. T., Suite of activity-based probes for cellulose-degrading enzymes. J Am Chem Soc 2012, 134 (50), 20521–32.

37. Lu, C. P.; Ren, C. T.; Lai, Y. N.; Wu, S. H.; Wang, W. M.; Chen, J. Y.; Lo, L. C., Design of a mechanism-based probe for neuraminidase to capture influenza viruses. Angew Chem Int Ed Engl 2005, 44 (42), 6888–92.

38. Lumba, M. A.; Willis, L. M.; Santra, S.; Rana, R.; Schito, L.; Rey, S.; Wouters, B. G.; Nitz, M., A β-galactosidase probe for the detection of cellular senescence by mass cytometry. Organic & Biomolecular Chemistry 2017, 15 (30), 6388–6392.

39. Tsai, C.-S.; Li, Y.-K.; Lo, L.-C., Design and synthesis of activity probes for glycosidases. Organic Letters 2002, 4 (21), 3607–3610.

40. Jiang, J.; Tan, Q.; Zhao, S.; Song, H.; Hu, L.; Xie, H., Late-stage difluoromethylation leading to a self-immobilizing fluorogenic probe for the visualization of enzyme activities in live cells. Chem Commun 2019, 55 (99), 15000–15003.

41. Gao, Z.; Thompson, A. J.; Paulson, J. C.; Withers, S. G., Proximity ligation-based fluorogenic imaging agents for neuraminidases. Angew. Chem. Int. Ed. 2018, 57(41), 13538–13541.

42. Huang, Y.-Y.; Kuo, C.-C.; Chu, C.-Y.; Huang, Y.-H.; Hu, Y.-L.; Lin, J.-J.; Lo, L.-C., Development of activity-based probes with tunable specificity for protein tyrosine phosphatase subfamilies. Tetrahedron 2010, 66 (25), 4521–4529.

43. Tai, C. H.; Lu, C. P.; Wu, S. H.; Lo, L. C., Synthesis and evaluation of turn-on fluorescent probes for imaging steroid sulfatase activities in cells. Chemical communications 2014, 50 (46), 6116–9.

44. Mao, W.; Xia, L.; Wang, Y.; Xie, H., A Self-Immobilizing and Fluorogenic Probe for beta-Lactamase Detection. Chem. Asian J. 2016, 11 (24), 3493–3497.

45. Doura, T.; Kamiya, M.; Obata, F.; Yamaguchi, Y.; Hiyama, T. Y.; Matsuda, T.; Fukamizu, A.; Noda, M.; Miura, M.; Urano, Y., Detection of lacZ-positive cells in living tissue with single-cell resolution. Angew. Chem. Int. Ed. 2016, 55 (33), 9620–4.

46. Ito, H.; Kawamata, Y.; Kamiya, M.; Tsuda-Sakurai, K.; Tanaka, S.; Ueno, T.; Komatsu, T.; Hanaoka, K.; Okabe, S.; Miura, M.; Urano, Y., Red-shifted fluorogenic substrate for detection of lacZ-positive cells in living tissue with single-cell resolution. Angew. Chem. Int. Ed. 2018, 57 (48), 15702–15706.

47. Liu, J.; Cai, L.; Sun, W.; Cheng, R.; Wang, N.; Jin, L.; Rozovsky, S.; Seiple, I. B.; Wang, L., Photocaged quinone methide crosslinkers for light-controlled chemical crosslinking of protein-protein and protein-DNA complexes. Angew. Chem. Int. Ed. 2019, 58 (52), 18839–18843.

48. Ahmed, V.; Liu, Y.; Taylor, S. D., Multiple pathways for the irreversible inhibition of steroid sulfatase with quinone methide-generating suicide inhibitors. ChemBioChem 2009, 10 (9), 1457–1461.

49. Liu, J.; Ma, X.; Cui, C.; Wang, Y.; Deenik, P. R.; Cui, L., A self-immobilizing NIR probe for non-invasive imaging of senescence. BioRxiv 2020.

50. Li, Y.; Song, H.; Xue, C.; Fang, Z.; Xiong, L.; Xie, H., A self-immobilizing near-infrared fluorogenic probe for sensitive imaging of extracellular enzyme activity in vivo. Chemical Science 2020, 11 (23), 5889–5894.

51. Hyun, J. Y.; Park, S. H.; Park, C. W.; Kim, H. B.; Cho, J. W.; Shin, I., Trifunctional fluorogenic probes for fluorescence imaging and isolation of glycosidases in cells. Organic Letters 2019, 21 (12), 4439–4442.

