## Supplementary information for "A Versatile Linker for Probes Targeting Hydrolases via *In Situ* labeling"

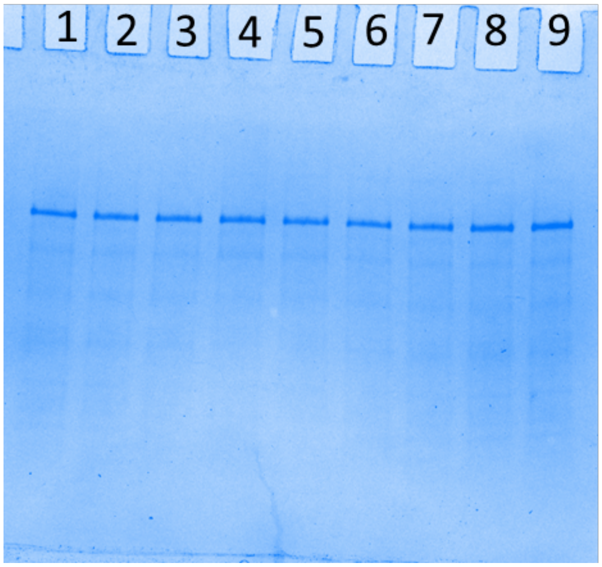


**Figure S1.** Coomassie staining of FITC-labeled β-Galactosidase (10 μg) after it was incubated with different concentrations of BG-FITC-2F, BG-FITC-0F or BG-FITC-1F. Lane 1-3: 1 μM of BG-FITC-2F, BG-FITC-0F or BG-FITC-1F; Lane 4-6: 5 μM of BG-FITC-2F, BG-FITC-0F or BG-FITC-1F; Lane 7-8: 1 μM of BG-FITC-2F, BG-FITC-0F or BG-FITC-1F.

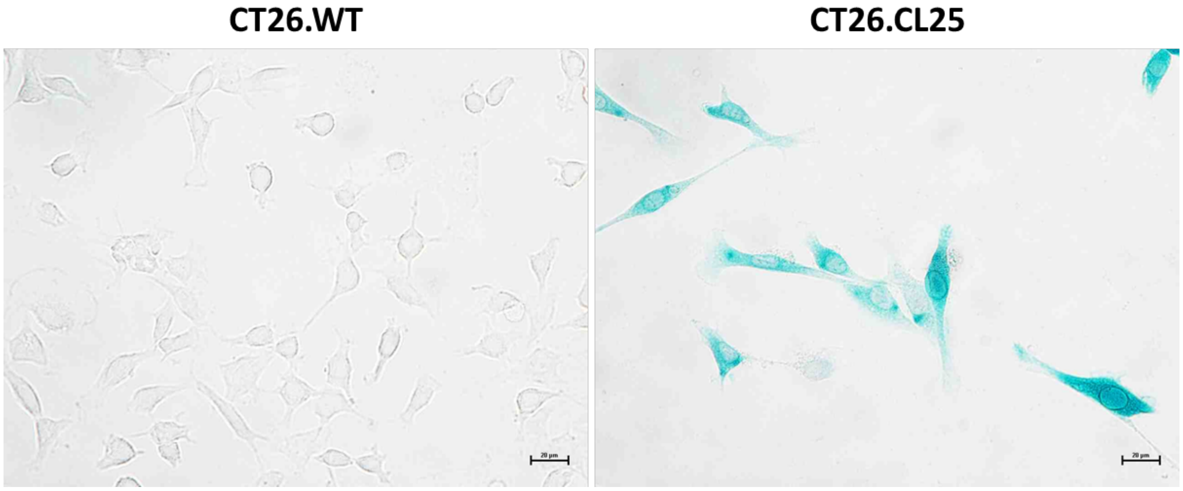


**Figure S2.** X-gal staining of CT26.WT and CT26.CL25. Scale bar: 20 μm.

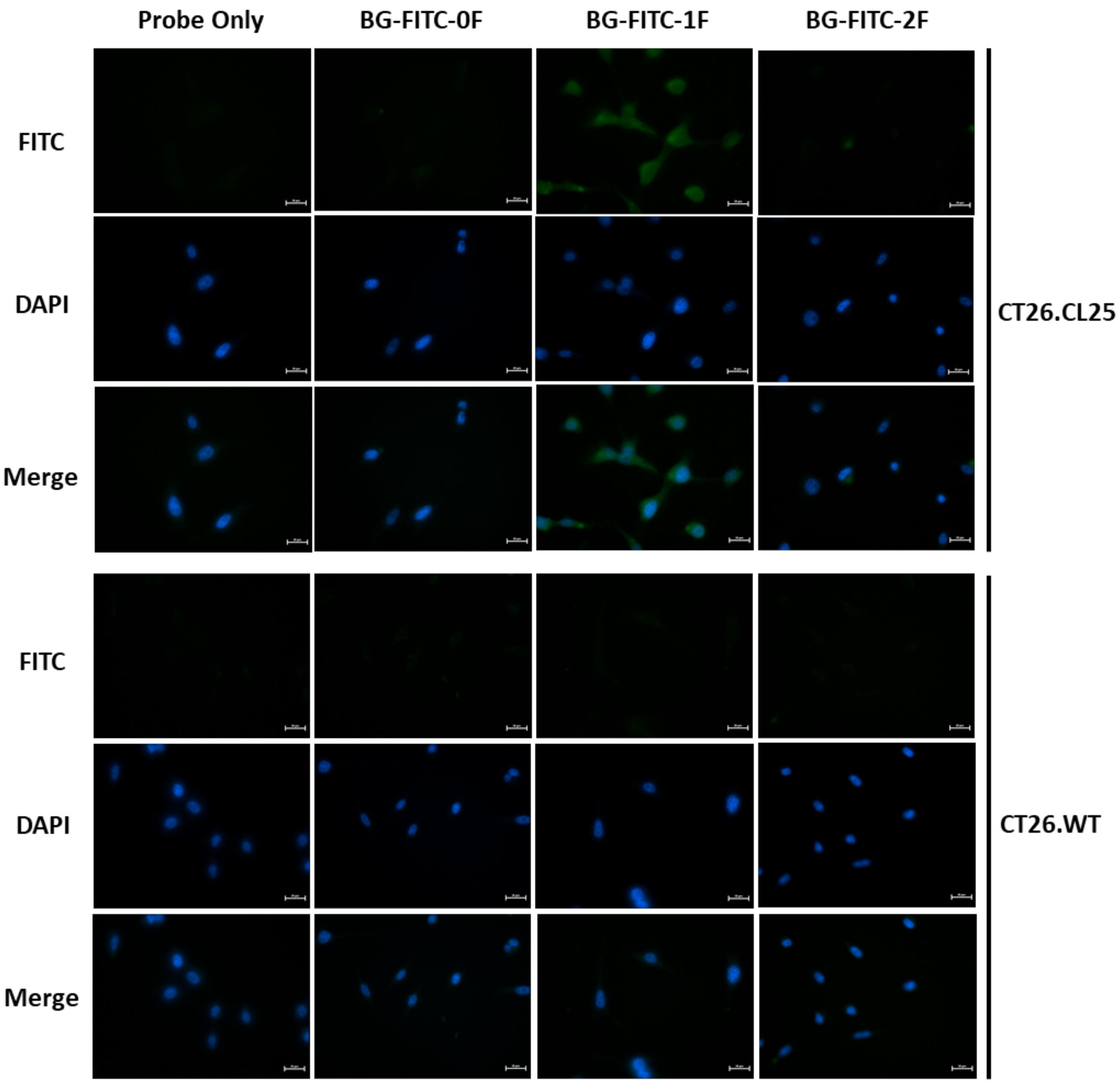


**Figure S3.** Fluorescent cell images of CT26.WT and CT26.CL25 cells after incubation with 10 μM of BG-FITC-0F, BG-FITC-1F or BG-FITC-2F for 14 hours. Scale bar: 20 μm.

### General Information for Chemical Synthesis

All chemicals were purchased from commercial sources unless otherwise noted. HRMS samples were analyzed on Waters LCT Premier Mass Spectrometer. High-performance liquid chromatography (HPLC) was performed on a Dionex Ultimate 300 HPLC System (Thermo Scientific) equipped with a GP50 gradient pump and an in-line diode array UV-Vis detector. Reverse-phase C18 columns were used with acetonitrile/water gradient mobile phase containing 0.1% trifluoroacetic acid. NMR spectra were recorded on Bruker instruments (500 MHz and 600 MHz for $^{1}$H NMR, 126 MHz and 151 MHz for $^{13}$C NMR) and internally referenced to the residual solvent signals ($^{1}$H: δ 7.26; $^{13}$C: δ 77.16 for $CDCl_3$, $^{1}$H: δ 3.31; $^{13}$C: δ 49.0 for $CD_3OD$ respectively). NMR chemical shifts (δ) and the coupling constants (J) for $^{1}$H and $^{13}$C NMR are reported in parts per million (ppm) and in Hertz, respectively. The following conventions are used for multiplicities: s, singlet; d, doublet; t, triplet; m, multiplet; and dd, doublet of doublet. Compounds **1**, **2**, **5** were prepared following the literature procedures(Chauvigne-Hines et al., 2012;Lilley et al., 2020).

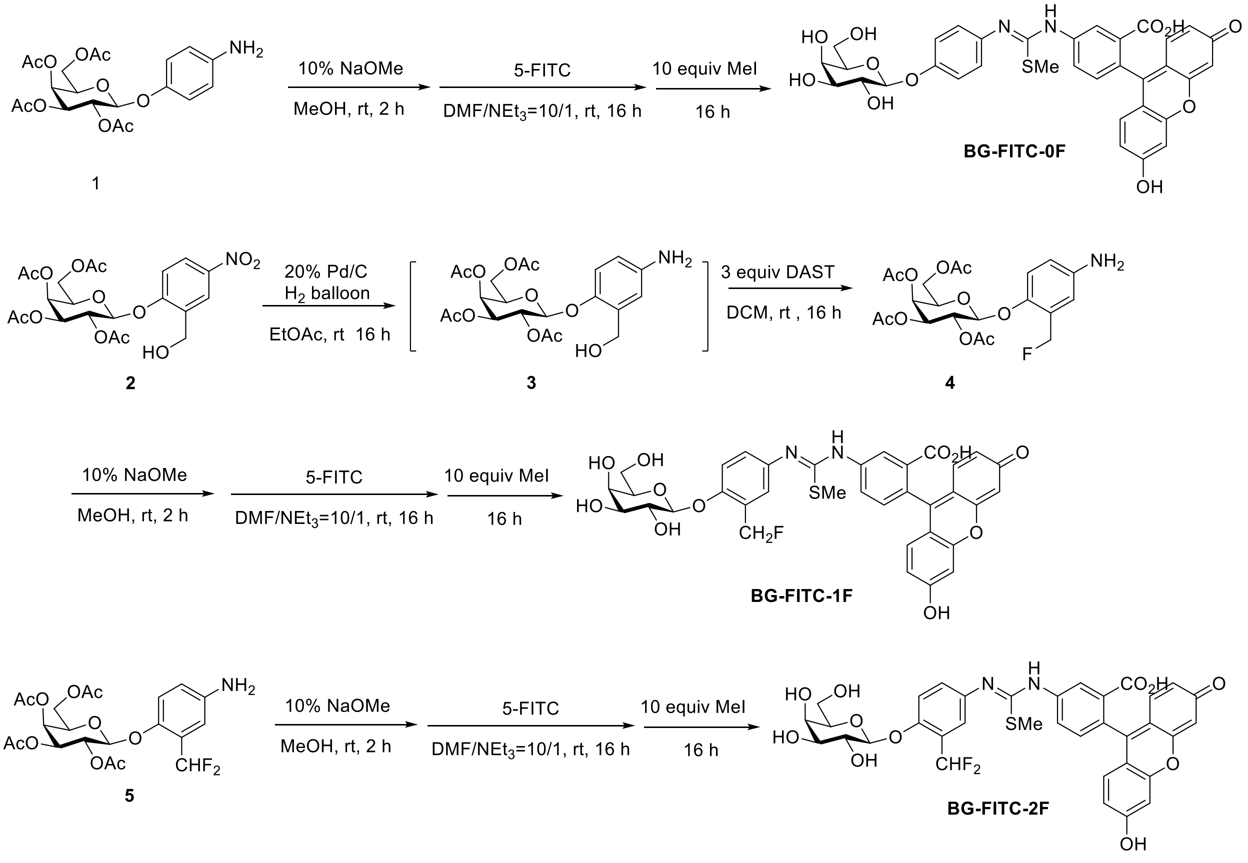


**Scheme S1.** Synthesis of BG-FITC-0F, BG-FITC-1F and BG-FITC-2F.

**1-(2-hydroxymethyl-4-aminophenyl)-2,3,4,6-tetra-*O*-acetyl-β-D-galactopyranose (3)**: To a solution of compound **2** in EtOAc was added 20% wt Pd/C on carbon. Then the reaction mixture was purged with H2 balloon three times and was stirred at room temperature for 16 hours. The mixture was filtered, and the filtrate concentrated by rotary evaporation and further purified by column chromatography to afford compound **3**, which was used in the next step without further purifications.

**1-(2-fluoromethyl-4-aminophenyl)-2,3,4,6-tetra-*O*-acetyl-β-D-galactopyranose (4)**: To a solution of compound 3 in DCM was added 3 equiv. DAST. Then the reaction mixture was stirred at room temperature for 16 hours. The reaction mixture was quenched by the addition of water and extracted with DCM and washed with brine. The organic layer was dried over Na2SO4, filtered and concentrated by rotary evaporation. The residue was purified by column chromatography to afford compound **4.** Yield: 70% over 2 steps.

**General procedure for deacetylation, conjugation with 5-FITC and methylation.**

The corresponding starting material was added to a 20 mL vial and methanol was added, followed by 10 mol% NaOMe. The mixture was stirred at room temperature until the reaction was complete monitored by HPLC. Upon completion, the reaction was quenched with acidic resin and filtered to give a crude product. This crude product was dissolved in a mixture of DMF/$NEt_3$ (10/1) and 5-FITC (1 equiv.) was added to the solution. The resulting mixture was stirred at room temperature monitored by HPLC. Upon completion, MeI (10 equiv.) was added to the above mixture and further stirred at room temperature for additional 16 hours. Then the reaction mixture was purified by semi-preparative HPLC. Yields: 50% for BG-FITC-0F, 50% for BG-FITC-1F and 73% for BG-FITC-2F.

**Compound 4**

**$^1$H NMR** (600 MHz, $CDCl_3$) δ 6.92 (dd, $J$ = 8.6, 0.9 Hz, 1H, Ar), 6.72 (d, $J$ = 2.0 Hz, 1H, Ar), 6.61 (ddd, $J$ = 8.7, 2.7, 1.6 Hz, 1H, Ar), 5.48 (dd, $J$ = 10.5, 8.0 Hz, 1H, H-2), 5.46 – 5.42 (m, 1H, H-4), 5.34 (dd, $J$ = 10.8, 126.6 Hz, 1H, C$H_2$F), 5.26 (dd, $J$ = 47.9, 11.0 Hz, 1H, C$H_2$F), 5.08 (dd, $J$ = 10.5, 3.4 Hz, 1H, H-3), 4.87 (d, $J$ = 8.0 Hz, 1H, H-1), 4.24 (dd, $J$ = 11.3, 7.0 Hz, 1H, H-6a),

4.15 (dd, *J* = 11.3, 6.4 Hz, 1H, H-6b), 4.04 – 3.98 (m, 1H, H-5), 3.74 – 3.34 (m, 2H, NH), 2.19 (s, 3H, OAc), 2.09 (s, 3H, OAc), 2.06 (s, 3H, OAc), 2.01 (s, 3H, OAc).

**$^{13}$C NMR** (151 MHz, $CDCl_3$) δ 170.51 (O*C*OCH3), 170.41 (O*C*OCH3), 170.28 (O*C*OCH3), 169.71 (O*C*OCH3), 147.55 (d, *J* = 4.8 Hz, Ar), 142.68, 127.61 (d, *J* = 16.6 Hz, Ar), 117.85 (Ar), 116.22 (d, *J* = 3.0 Hz, Ar), 115.96 (d, *J* = 7.4 Hz, Ar), 101.30 (C-1), 80.07 (d, *J* = 165.0 Hz, *C*H2F), 71.05 (C-5), 70.95 (C-3), 68.64 (C-2), 67.04 (C-4), 61.48 (C-6), 20.84 (OCO*C*H3), 20.82 (OCO*C*H3), 20.80 (OCO*C*H3), 20.73 (OCO*C*H3).

**$^{19}$F NMR** (565 MHz, $CDCl_3$) δ -212.87 (t, *J* = 47.6 Hz).

**LRMS (ESI)**: calc'd for $C_{21}H_{27}FNO_{10}^+$ [$(M+H)^+$] 472.15; found: 472.13.

**Compound 5**

**$^{1}$H NMR** (600 MHz, $CDCl_3$) δ 6.94 (d, *J* = 8.7 Hz, 1H, Ar), 6.86 (dd, *J* = 2.9, 1.4 Hz, 1H, Ar), 6.74 (d, *J* = 54 Hz, C*H*F2) 6.72 – 6.67 (m, 1H, Ar), 5.47 (dd, *J* = 10.5, 8.0 Hz, 1H, H-2), 5.44 (dd, *J* = 3.5, 1.1 Hz, 1H, H-4), 5.08 (dd, *J* = 10.5, 3.5 Hz, 1H, H-3), 4.86 (d, *J* = 8.0 Hz, 1H, H-1), 4.23 (dd, *J* = 11.3, 7.0 Hz, 1H, H-6a), 4.14 (dd, *J* = 11.3, 6.4 Hz, 1H, H-6b), 4.02 (td, *J* = 6.7, 1.2 Hz, 1H, H-5), 3.87 – 3.47 (b, s, 2H, NH), 2.18 (s, 3H, OAc), 2.07 (s, 3H, OAc), 2.05 (s, 3H, OAc), 2.00 (s, 3H, OAc).

**$^{13}$C NMR** (151 MHz, $CDCl_3$) δ 170.36 (O*C*OCH3), 170.24 (O*C*OCH3), 170.12 (O*C*OCH3), 169.63 (O*C*OCH3), 147.24 (d, *J* = 6 Hz, Ar), 142.70 (Ar), 125.47 (t, *J* = 22.7 Hz, *C*H2F), 125.33 (Ar), 125.17 (Ar), 118.02 (Ar), 117.86 (Ar), 101.09 (C-1), 71.00 (C-5), 70.71 (C-3), 68.36 (C-2), 66.85 (C-4), 61.33 (C-6), 20.66 (OCO*C*H3), 20.64(OCO*C*H3), 20.60 (OCO*C*H3), 20.58 (OCO*C*H3).

**$^{19}$F NMR** (565 MHz, $CDCl_3$) δ -108.38 (dd, *J* = 300.9, 55.9 Hz), -121.57 (dd, *J* = 300.9, 54.9 Hz).

**LRMS (ESI)**: calc'd for $C_{21}H_{27}FNO_{10}^+$ [$(M+H)^+$]

**BG-FITC-1F**

**$^{1}$H NMR** (600 MHz, $CD_3OD$) δ 8.02 (s, 1H, Ar), 7.75 (d, *J* = 6.9 Hz, 1H, Ar), 7.47 – 7.17 (m, 4H, Ar), 6.76 (s, 2H, Ar), 6.66 (s, 4H, Ar), 5.64 – 5.46 (m, 2H, C*H*$_2$F), 4.92 (d, *J* = 7.8 Hz, 1H, H-1), 3.94 (d, *J* = 3.1 Hz, 1H, H-4), 3.87 (dd, *J* = 9.7, 7.8 Hz, 1H, H-2), 3.78 – 3.74 (m, 2H,H-6), 3.74 – 3.67 (m, 1H, H-5), 3.60 (dd, *J* = 9.7, 3.4 Hz, 1H, H-3), 2.82 (s, 3H, -SC*H*$_3$).

**$^{13}$C NMR** (151 MHz, $CD_3OD$) δ 169.93 (*C*=O), 162.15 (*C*OOH), 161.91 (S*C*(=N)N), 155.87 (Ar), 154.45 (Ar), 139.47 (Ar), 131.42 (Ar), 130.37 (Ar), 130.34 (Ar), 130.01 (Ar), 129.60 (Ar), 129.48 (Ar), 127.30 (Ar), 126.33 (Ar), 124.88 (Ar), 122.74 (Ar), 118.70 (Ar), 117.88 (Ar), 117.73 (Ar), 116.77 (Ar), 115.14 (Ar), 114.28 (Ar), 111.16 (Ar), 103.60 (Ar), 103.40 (Ar), 103.37 (Ar), 103.24 (C-1), 80.48 (d, *J* = 166 Hz, *C*H$_2$F), 77.23 (C-5), 74.87 (C-3), 72.13 (C-2), 70.19 (C-4), 62.42 (C-6), 15.11 (-S*C*H$_3$) .

**HRMS (ESI)**: calc'd for $C_{35}H_{30}FN_2O_{11}S^-$ [(M-H)$^-$] 705.1560; found: 705.1562.

**HPLC:**

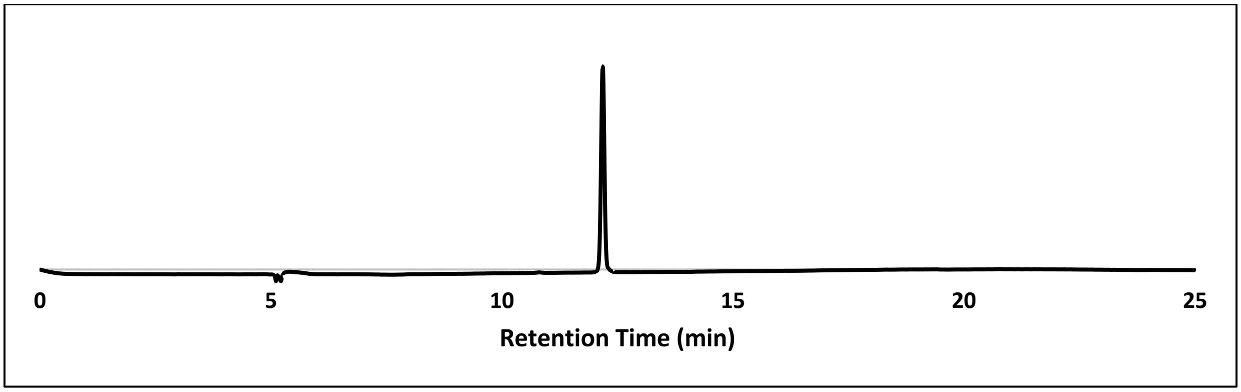


**BG-FITC-2F**

**$^{1}$H NMR** (600 MHz, $CD_3OD$) δ 8.02 (s, 1H, Ar), 7.74 (d, *J* = 6.0 Hz, 1H, Ar), 7.61 (s, 1H, Ar), 7.51 (d, *J* = 7.2 Hz, 1H, Ar), 7.43 (d, *J* = 8.9 Hz, 1H, Ar), 7.33 – 7.26 (m, 1H, Ar), 7.18 (t, *J* = 55.2 Hz, 1H, C*H*F$_2$), 6.76 (s, 2H, Ar), 6.66 (s, 4H, Ar), 4.94 (d, *J* = 7.8 Hz, 1H, H-1), 3.94 (d, *J* = 3.3 Hz, 1H, H-4), 3.88 (dd, *J* = 9.7, 7.8 Hz, 1H, H-2), 3.79 – 3.75 (m, 2H, H-6), 3.74 – 3.71 (m, 1H, H-5), 3.61 (dd, *J* = 9.7, 3.4 Hz, 1H, H-3), 2.79 (s, 3H, -SC*H*$_3$).

**$^{13}$C NMR** (151 MHz, $CD_3OD$) δ 169.92 (*C*=O), 161.72 (*C*OOH), 161.47 (S*C*(=N)N), 156.35 (Ar), 154.66 (Ar), 133.07 (Ar), 132.35 (Ar), 130.50 (Ar), 130.16 (Ar), 127.50 (Ar), 126.66 (Ar), 126.34 (Ar), 124.32 (Ar), 123.01 (Ar), 118.70 (Ar), 118.48 (Ar), 116.56 (Ar), 114.65 (Ar),

114.46 (Ar), 112.11 (t, *J* = 235 Hz), 111.42 (Ar), 103.68 (C-1), 103.59 (Ar) , 77.35 (C-5), 74.80 (C-3), 72.11 (C-2), 70.18 (C-4), 62.41 (C-6), 15.12 (-S*C*$H_3$).

**HRMS (ESI)**: calc'd for $C_{35}H_{31}F_2N_2O_{11}S^+$ [$(M+H)^+$] 725.1611; found: 725.1628.

**HPLC:**

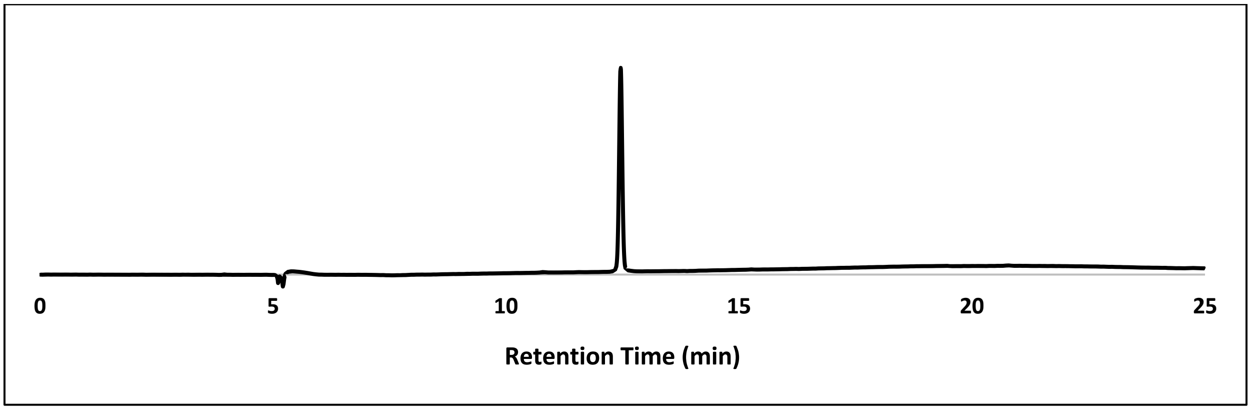


**BG-FITC-0F**

**$^1$H NMR** (500 MHz, $CD_3OD$) δ 8.04 (s, 1H, Ar), 7.76 (d, *J* = 7.8 Hz, 1H, Ar), 7.32 (d, *J* = 8.0 Hz, 3H, Ar), 7.19 (d, *J* = 8.5 Hz, 2H, Ar), 6.80 (s, 2H, Ar), 6.73 (d, *J* = 12.9 Hz, 4H, Ar), 4.89 (d, *J* = 7.7 Hz, 1H, H-1), 3.92 (d, *J* = 3.1 Hz, 1H, H-4), 3.88 – 3.79 (m, 1H, H-2), 3.75 – 3.70 (m, 3H, H-6, H-5), 3.60 (dd, *J* = 9.7, 3.2 Hz, 1H, H-3), 2.80 (s, 3H, -SC$H_3$).

**$^{13}$C NMR** (126 MHz, $CD_3OD$) δ 169.42 (*C*=O), 163.47 (*C*OOH), 162.20 (S*C*(=N)N), 161.90 (Ar), 161.61 (Ar), 161.32 (Ar), 159.42 (Ar), 155.23 (Ar), 138.77 (Ar), 133.40 (Ar), 130.87 (Ar), 130.80 (Ar), 130.60 (Ar), 130.51 (Ar), 128.40 (Ar), 128.08 (Ar), 124.40 (Ar), 121.01 (Ar), 118.94 (Ar), 118.71 (Ar), 116.40 (Ar), 115.23 (Ar), 111.96 (Ar), 103.58 (Ar), 102.86 (C-1), 77.06 (C-5), 74.74 (C-3), 72.16 (C-2), 70.15 (C-4), 62.38 (C-6), 15.09 (-S*C*$H_3$).

**HRMS (ESI)**: calc'd for $C_{34}H_{29}N_2O_{11}S^-$ [$(M-H)^-$] 673.1498; found: 673.1481.

**HPLC:**

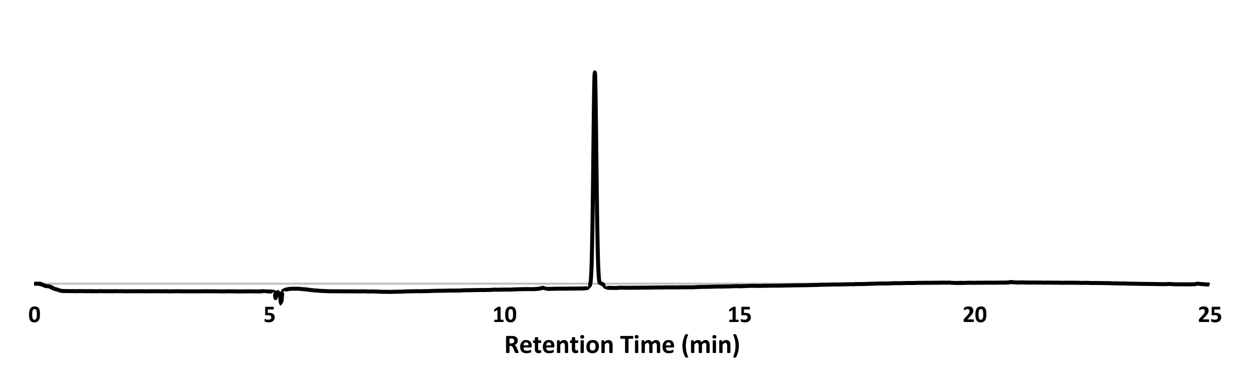


**Referrence:**


Chauvigne-Hines, L.M., Anderson, L.N., Weaver, H.M., Brown, J.N., Koech, P.K., Nicora, C.D., Hofstad, B.A., Smith, R.D., Wilkins, M.J., Callister, S.J., and Wright, A.T. (2012). Suite of activity-based probes for cellulose-degrading enzymes. *J Am Chem Soc* 134**,** 20521-20532.

Lilley, L.M., Kamper, S., Caldwell, M., Chia, Z.K., Ballweg, D., Vistain, L., Krimmel, J., Mills, T.A., Macrenaris, K., Lee, P., Waters, E.A., and Meade, T.J. (2020). Self-Immolative Activation of β-Galactosidase-Responsive Probes for In Vivo MR Imaging in Mouse Models. *Angewandte Chemie International Edition* 59**,** 388-394.


**NMR and Mass Spectra**

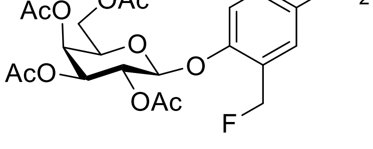


**4**

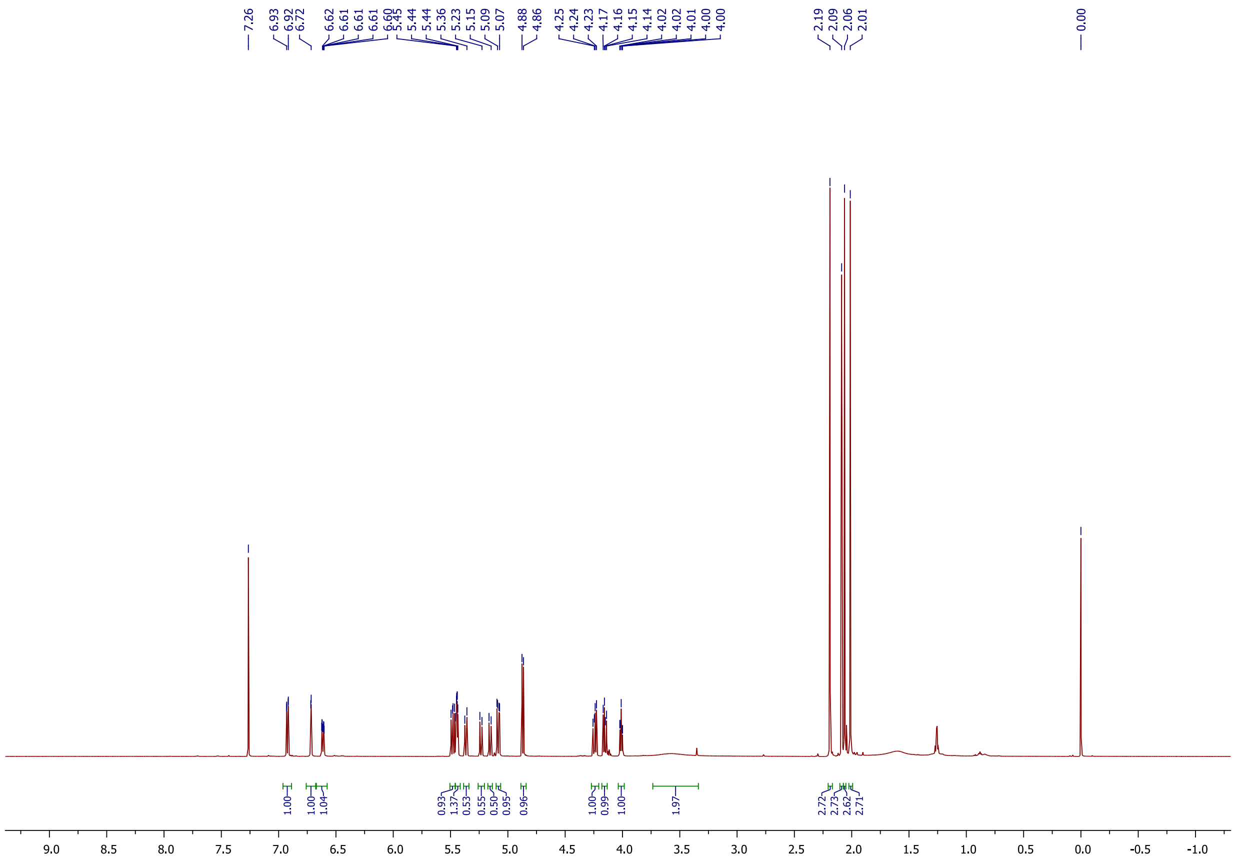

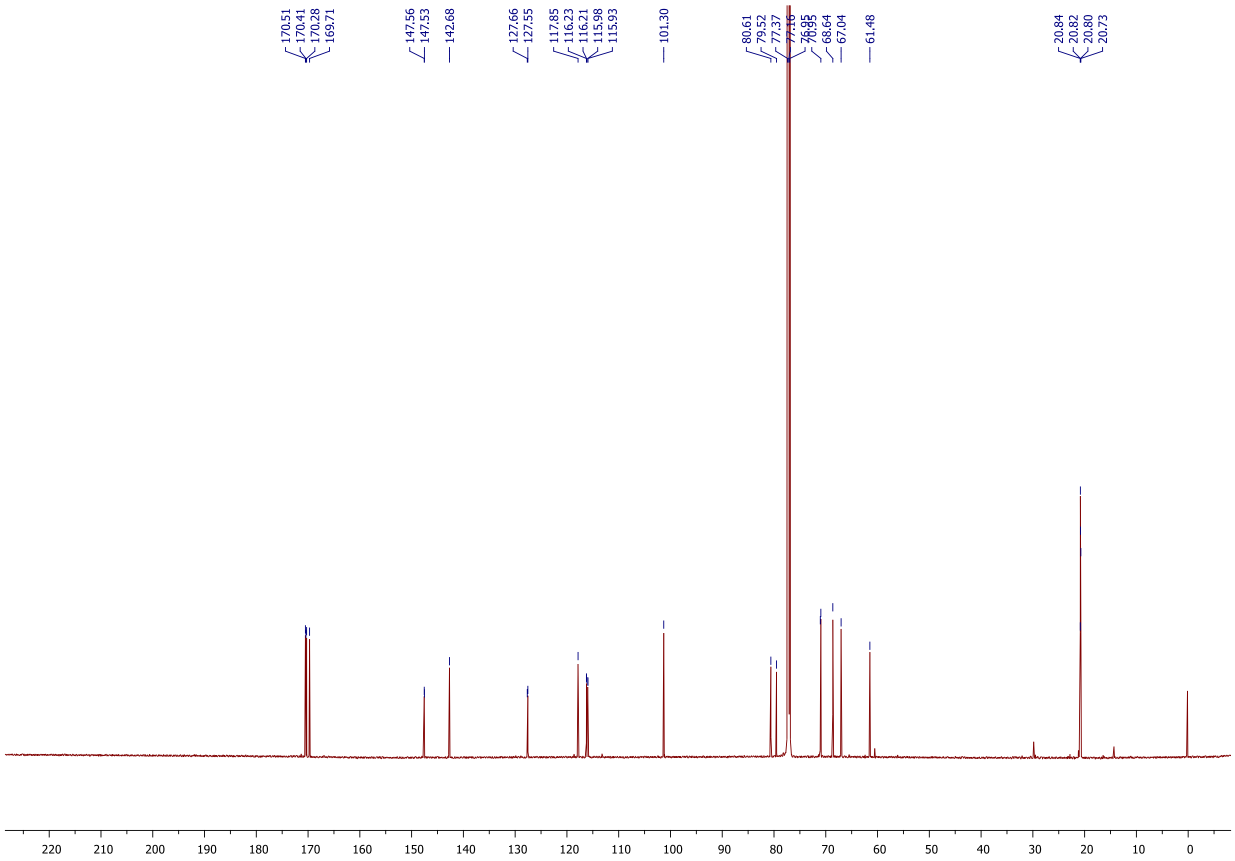
170.51
170.41
170.28
169.71
147.56
147.53
142.68
127.66
127.55
117.85
116.23
116.21
115.98
115.93
101.30
80.61
79.52
77.37
77.16
76.95
70.95
68.64
67.04
61.48
20.84
20.82
20.80
20.73
220 210 200 190 180 170 160 150 140 130 120 110 100 90 80 70 60 50 40 30 20 10 0

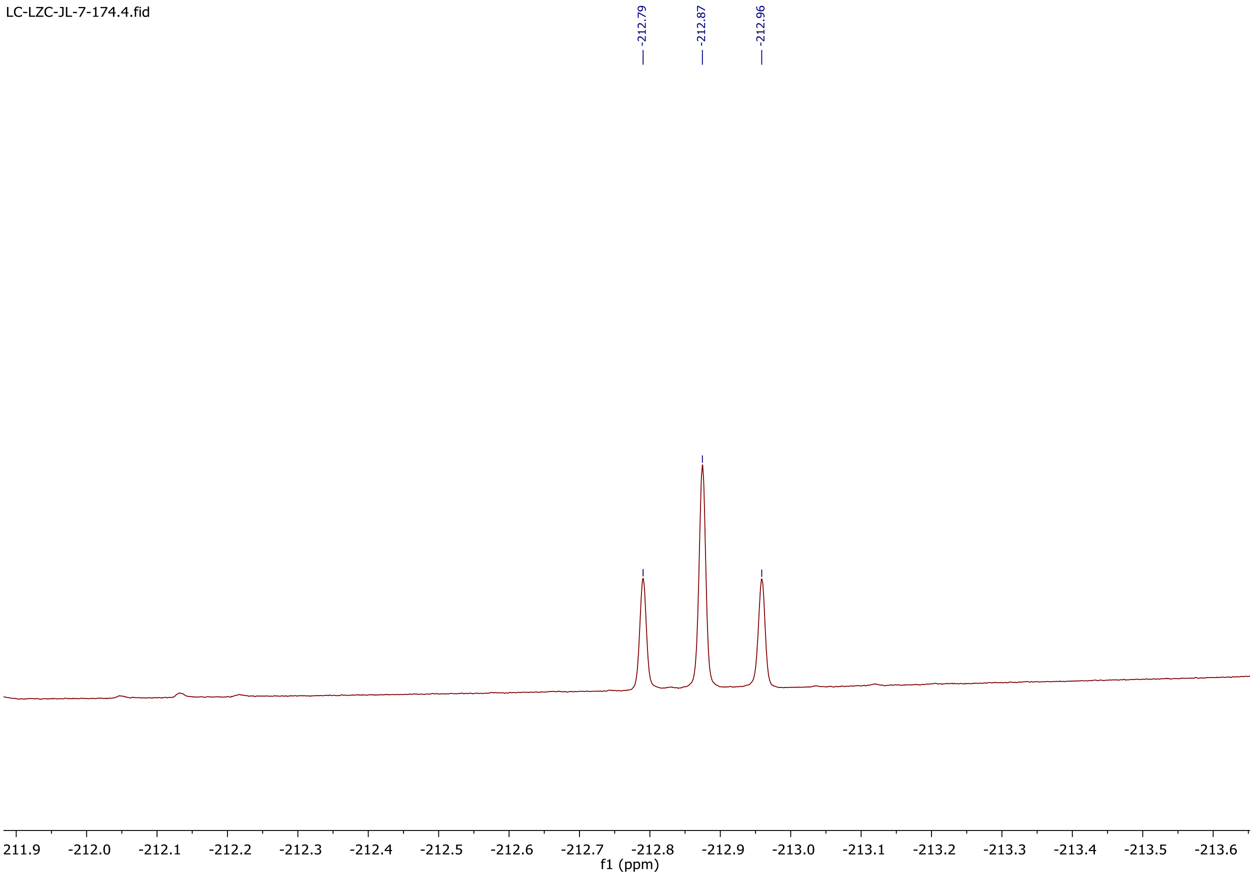
LC-LZC-JL-7-174.4.fid
-212.79
-212.87
-212.96
211.9
-212.0
-212.1
-212.2
-212.3
-212.4
-212.5
-212.6
-212.7
-212.8
-212.9
-213.0
-213.1
-213.2
-213.3
-213.4
-213.5
-213.6
f1 (ppm)

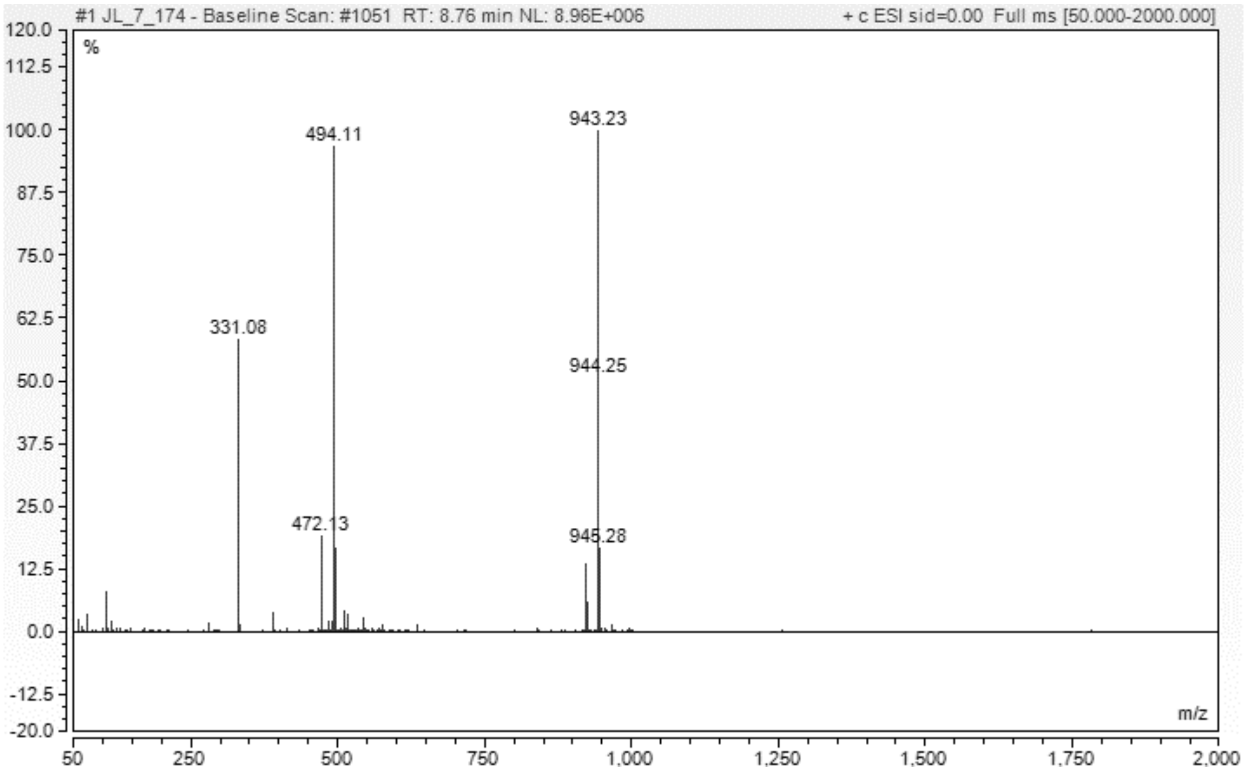
#1 JL_7_174 - Baseline Scan: #1051 RT: 8.76 min NL: 8.96E+006
+ c ESI sid=0.00 Full ms [50.000-2000.000]
%
943.23
494.11
331.08
944.25
472.13
945.28
m/z

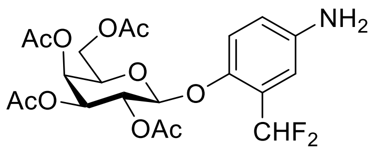

**5**

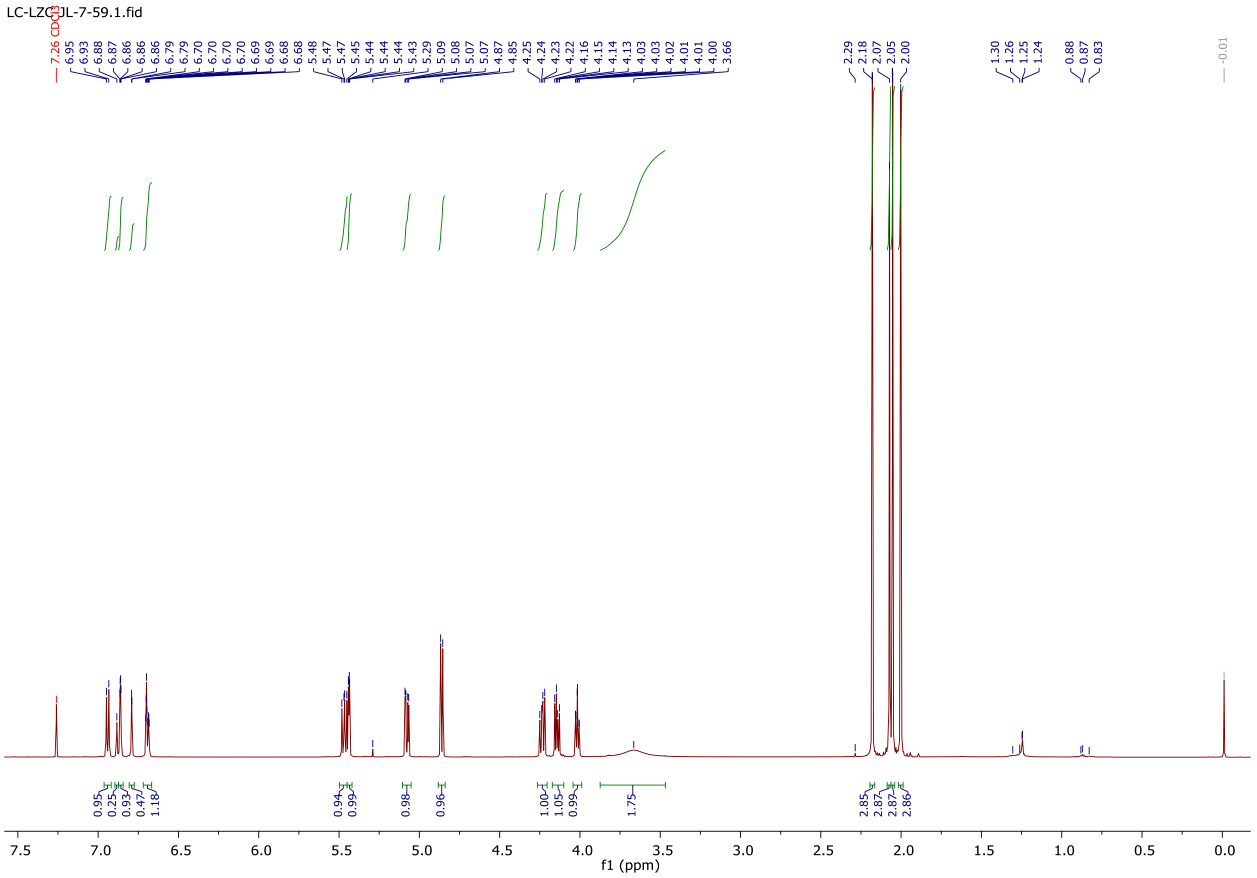

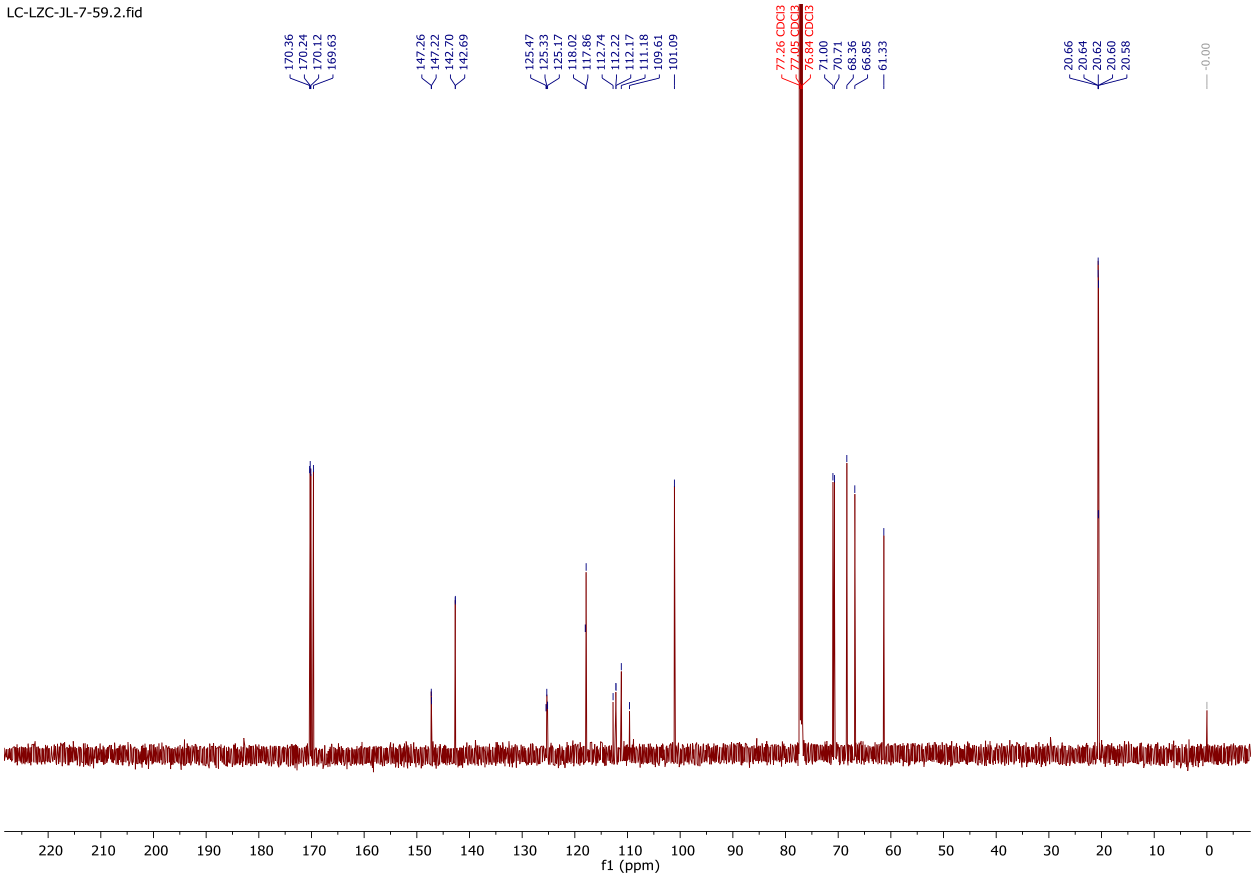
LC-LZC-JL-7-59.2.fid
170.36
170.24
170.12
169.63
147.26
147.22
142.70
142.69
125.47
125.33
125.17
118.02
117.86
112.74
112.22
112.17
111.18
109.61
101.09
77.26 CDCl3
77.05 CDCl3
76.84 CDCl3
71.00
70.71
68.36
66.85
61.33
20.66
20.64
20.62
20.60
20.58
-0.00
220
210
200
190
180
170
160
150
140
130
120
110
100
90
80
70
60
50
40
30
20
10
0
f1 (ppm)

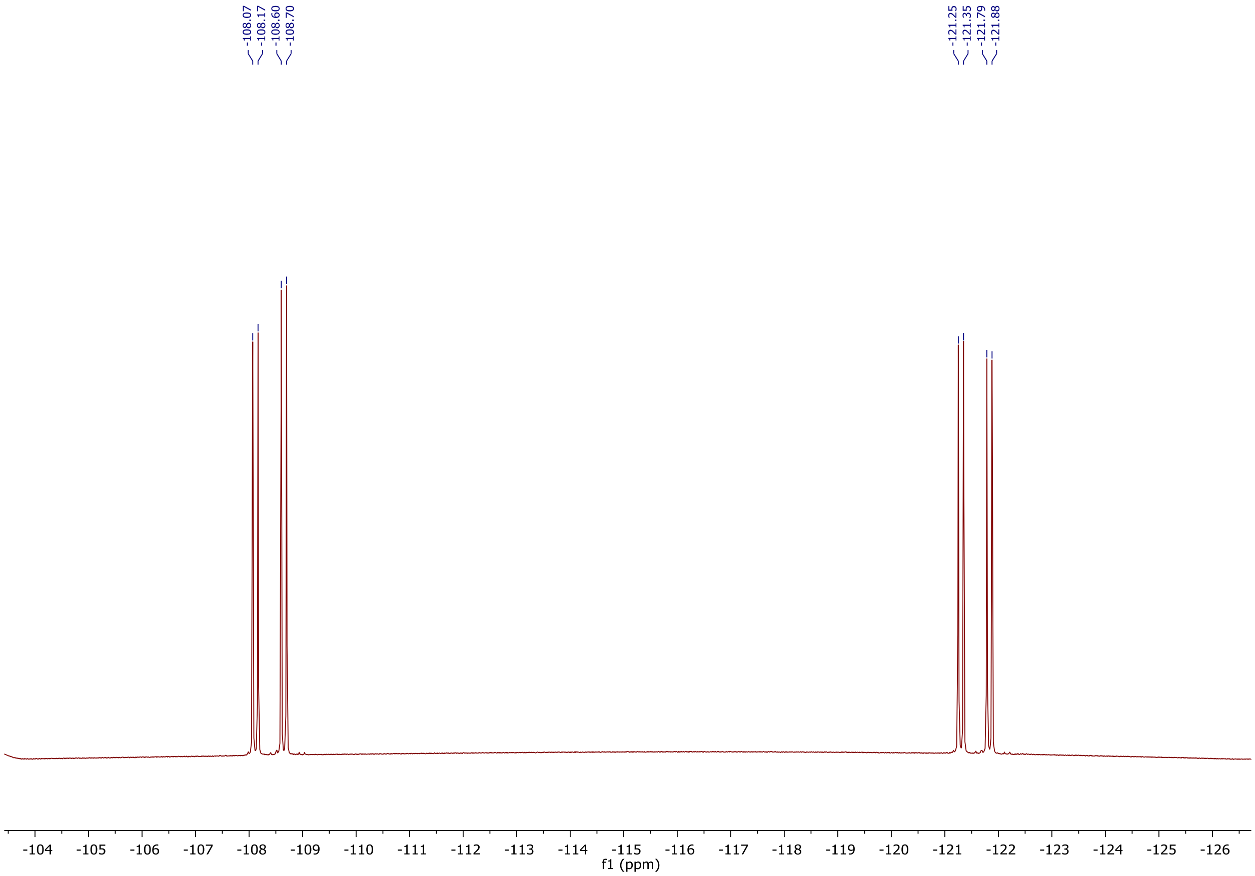
-108.07
-108.17
-108.60
-108.70
-121.25
-121.35
-121.79
-121.88
-104
-105
-106
-107
-108
-109
-110
-111
-112
-113
-114
-115
-116
-117
-118
-119
-120
-121
-122
-123
-124
-125
-126
f1 (ppm)

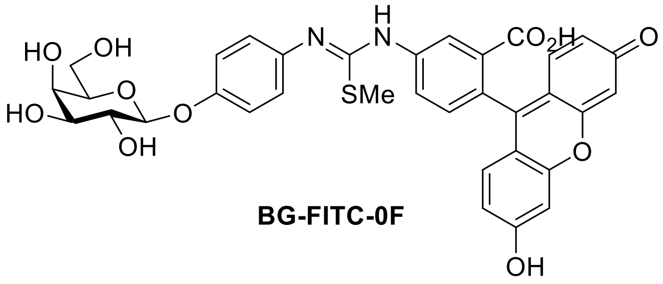

**BG-FITC-0F**

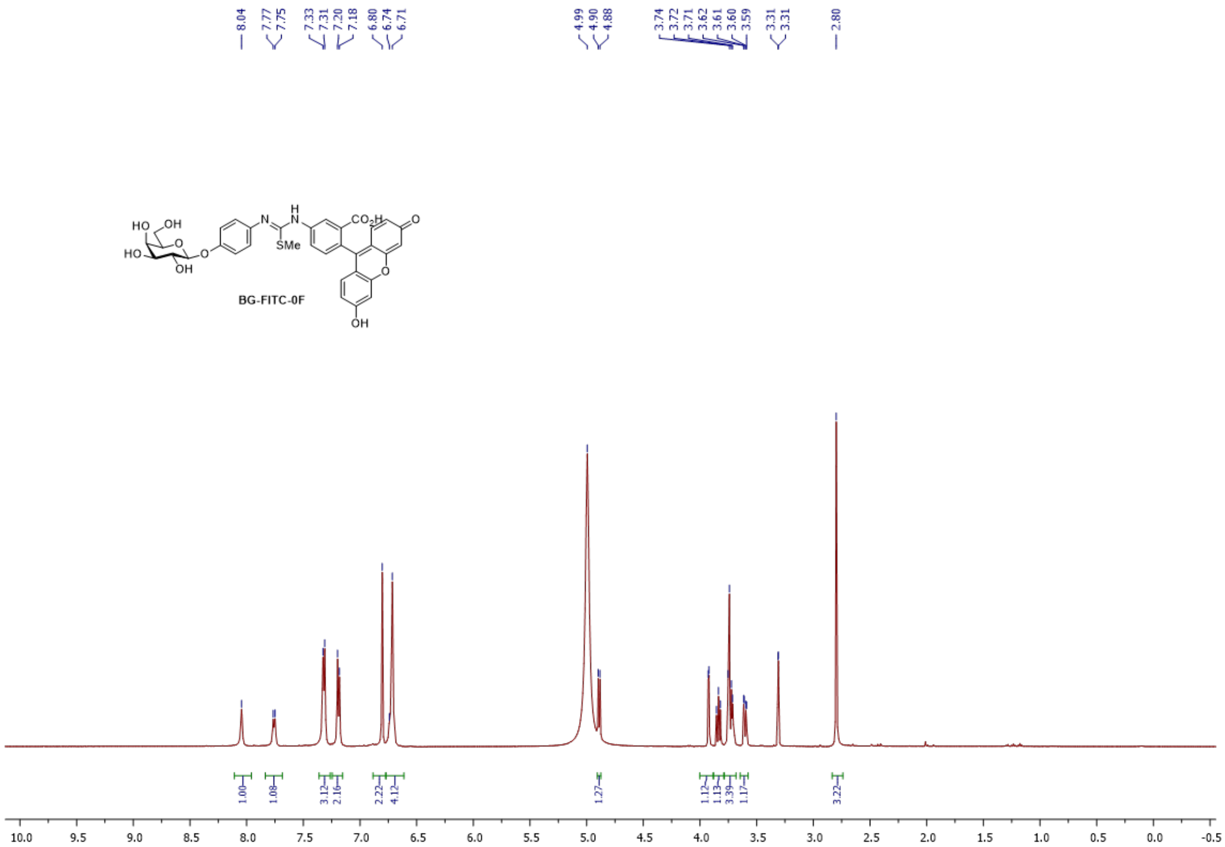
BG-FITC-0F
8.04
7.77
7.75
7.33
7.31
7.20
7.18
6.80
6.74
6.71
4.99
4.90
4.88
3.74
3.72
3.71
3.62
3.61
3.60
3.59
3.31
3.31
2.80
1.00
1.08
3.12
2.16
2.22
4.12
1.27
1.12
1.13
3.39
1.17
3.22
10.0 9.5 9.0 8.5 8.0 7.5 7.0 6.5 6.0 5.5 5.0 4.5 4.0 3.5 3.0 2.5 2.0 1.5 1.0 0.5 0.0 -0.5

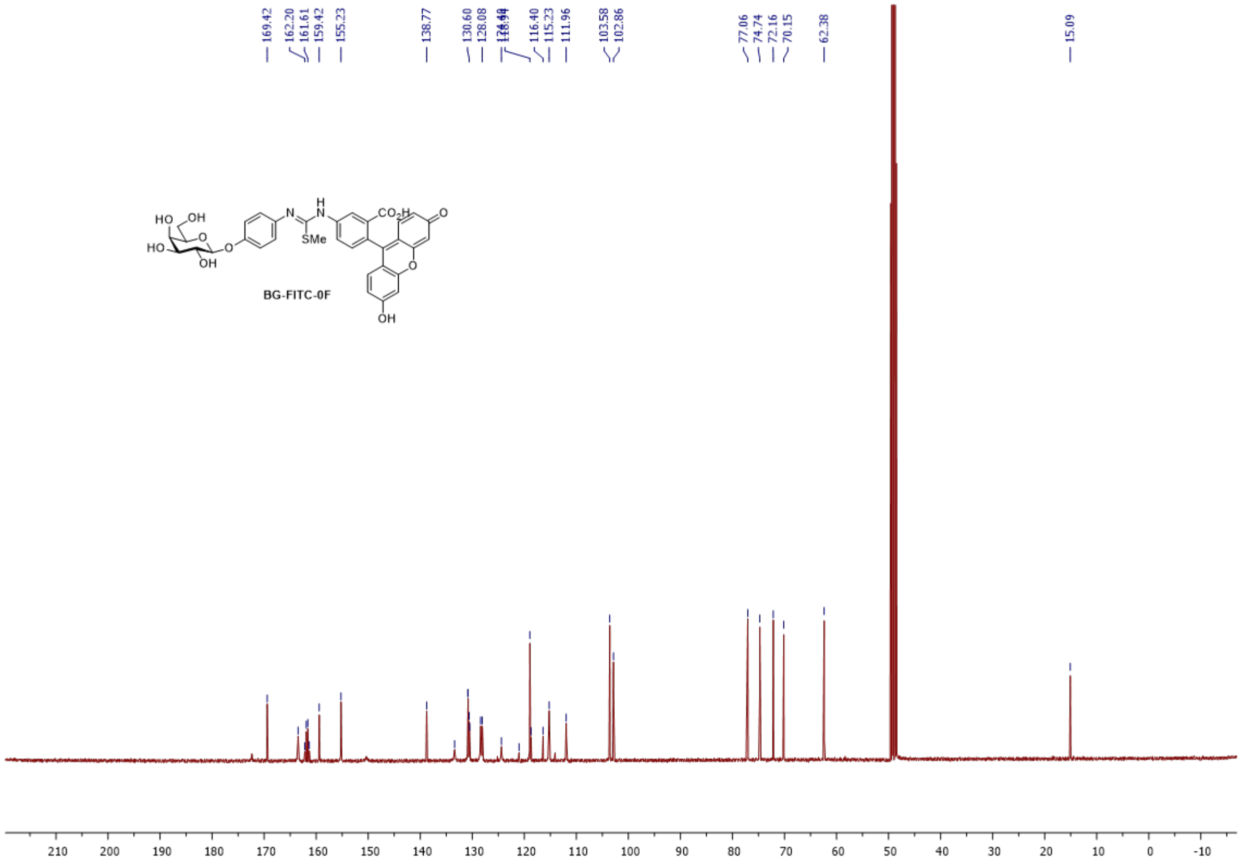
169.42
162.20
161.61
159.42
155.23
138.77
130.60
128.08
116.40
115.23
111.96
103.58
102.86
77.06
74.74
72.16
70.15
62.38
15.09
BG-FITC-0F
210 200 190 180 170 160 150 140 130 120 110 100 90 80 70 60 50 40 30 20 10 0 -10

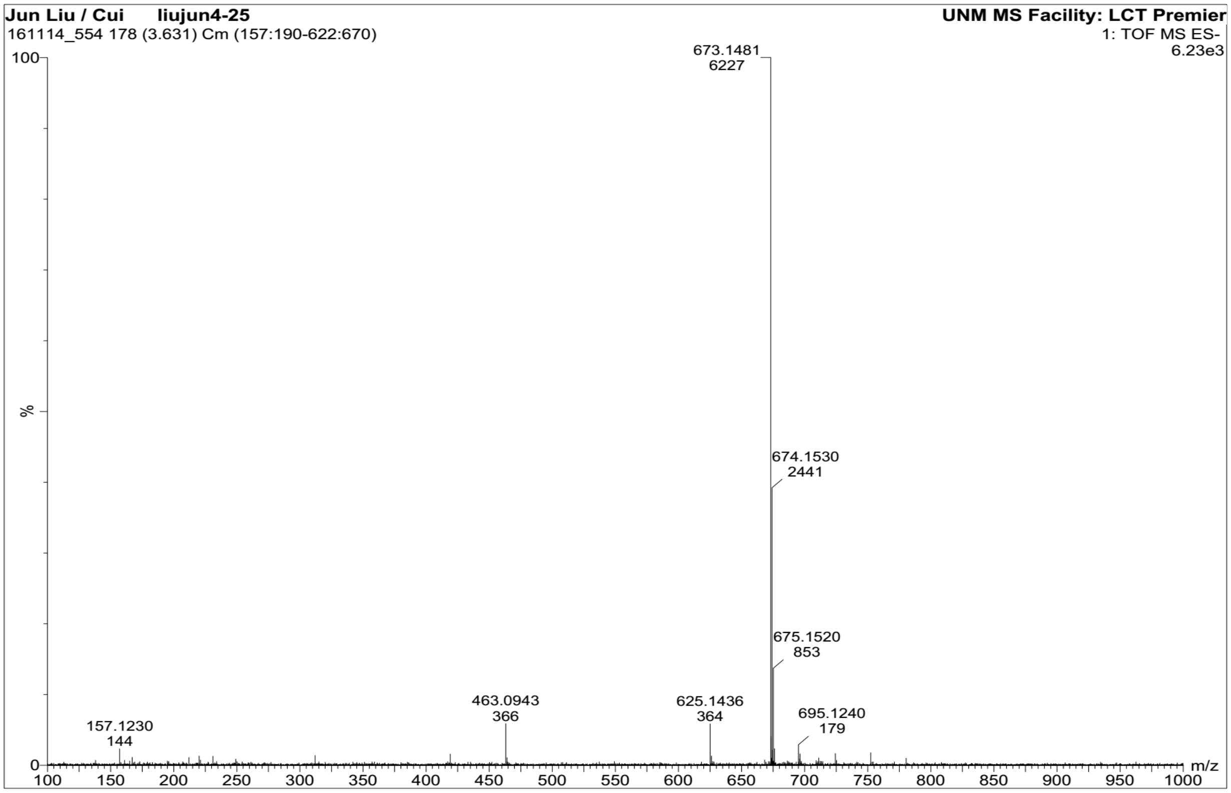
Jun Liu / Cui liujun4-25
161114_554 178 (3.631) Cm (157:190-622:670)
UNM MS Facility: LCT Premier
1: TOF MS ES-
6.23e3
673.1481
6227
674.1530
2441
675.1520
853
463.0943
366
625.1436
364
695.1240
179
157.1230
144
%
100
0
m/z
100 150 200 250 300 350 400 450 500 550 600 650 700 750 800 850 900 950 1000

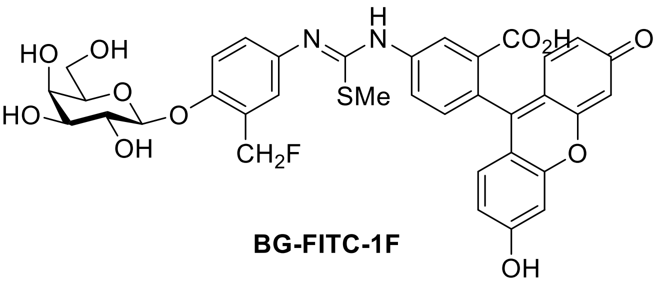

**BG-FITC-1F**

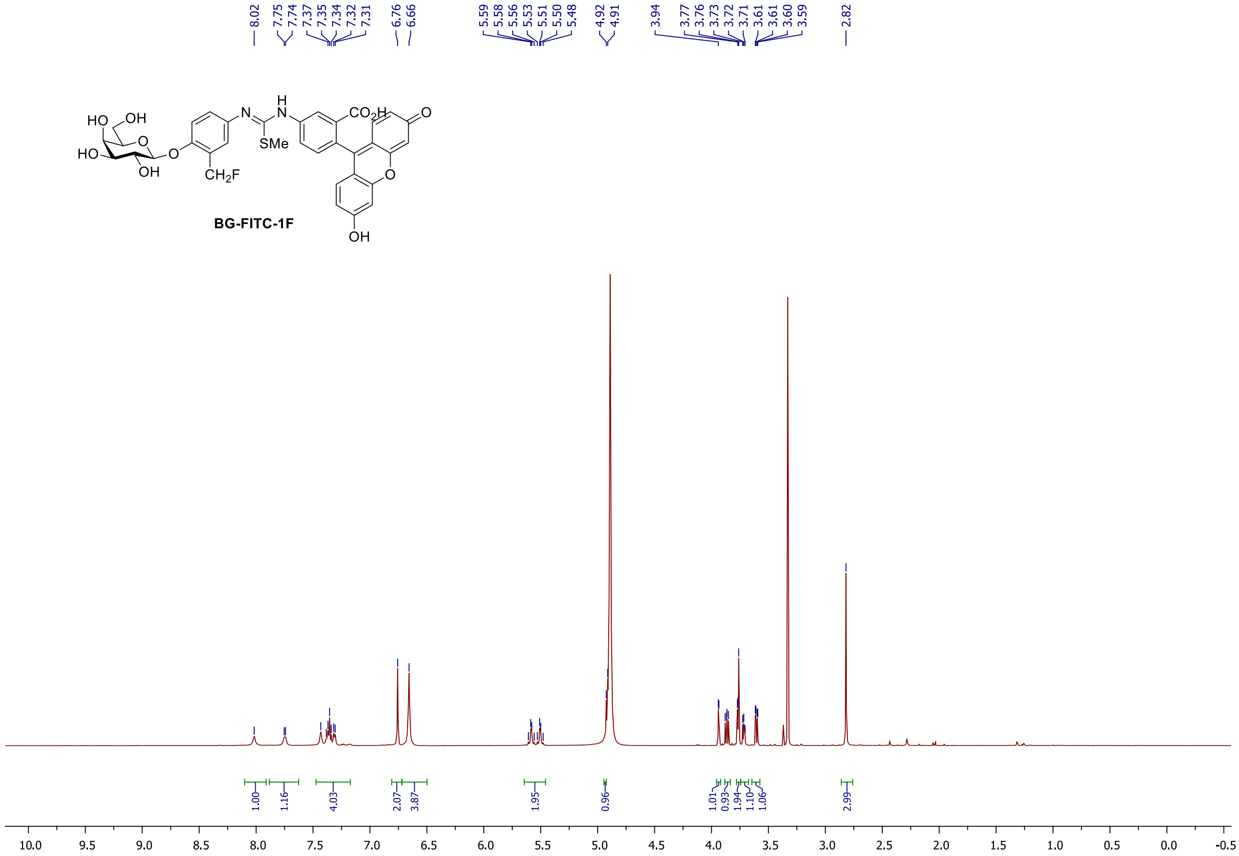
8.02
7.75 7.74 7.37 7.35 7.34 7.32 7.31
6.76 6.66
5.59 5.58 5.56 5.53 5.51 5.50 5.48
4.92 4.91
3.94 3.77 3.76 3.73 3.72 3.71 3.61 3.61 3.60 3.59
2.82
BG-FITC-1F
1.00 1.16 4.03 2.07 3.87 1.95 0.96 1.01 0.93 1.94 1.10 1.06 2.99
10.0 9.5 9.0 8.5 8.0 7.5 7.0 6.5 6.0 5.5 5.0 4.5 4.0 3.5 3.0 2.5 2.0 1.5 1.0 0.5 0.0 -0.5

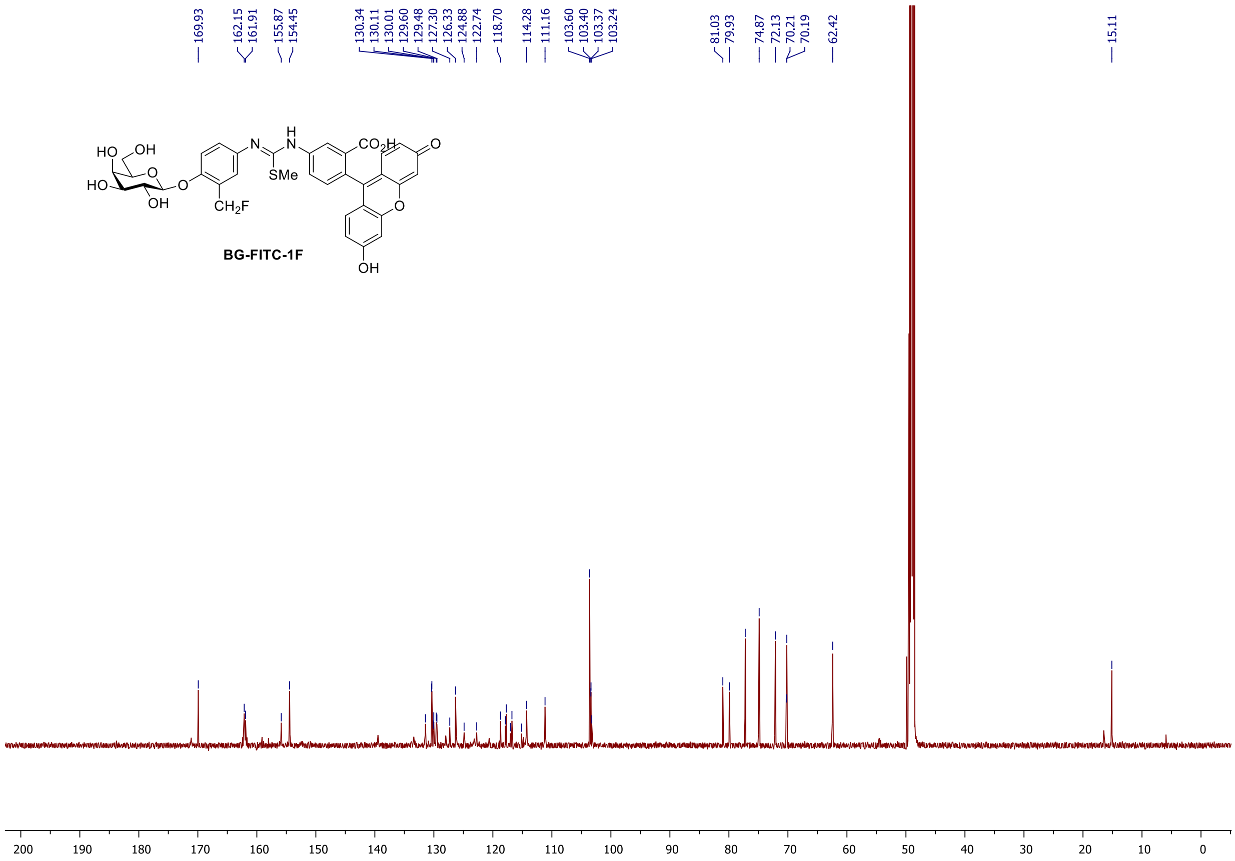
169.93
162.15
161.91
155.87
154.45
130.34
130.11
130.01
129.60
129.48
127.30
126.33
124.88
122.74
118.70
114.28
111.16
103.60
103.40
103.37
103.24
81.03
79.93
74.87
72.13
70.21
70.19
62.42
15.11
BG-FITC-1F
200 190 180 170 160 150 140 130 120 110 100 90 80 70 60 50 40 30 20 10 0

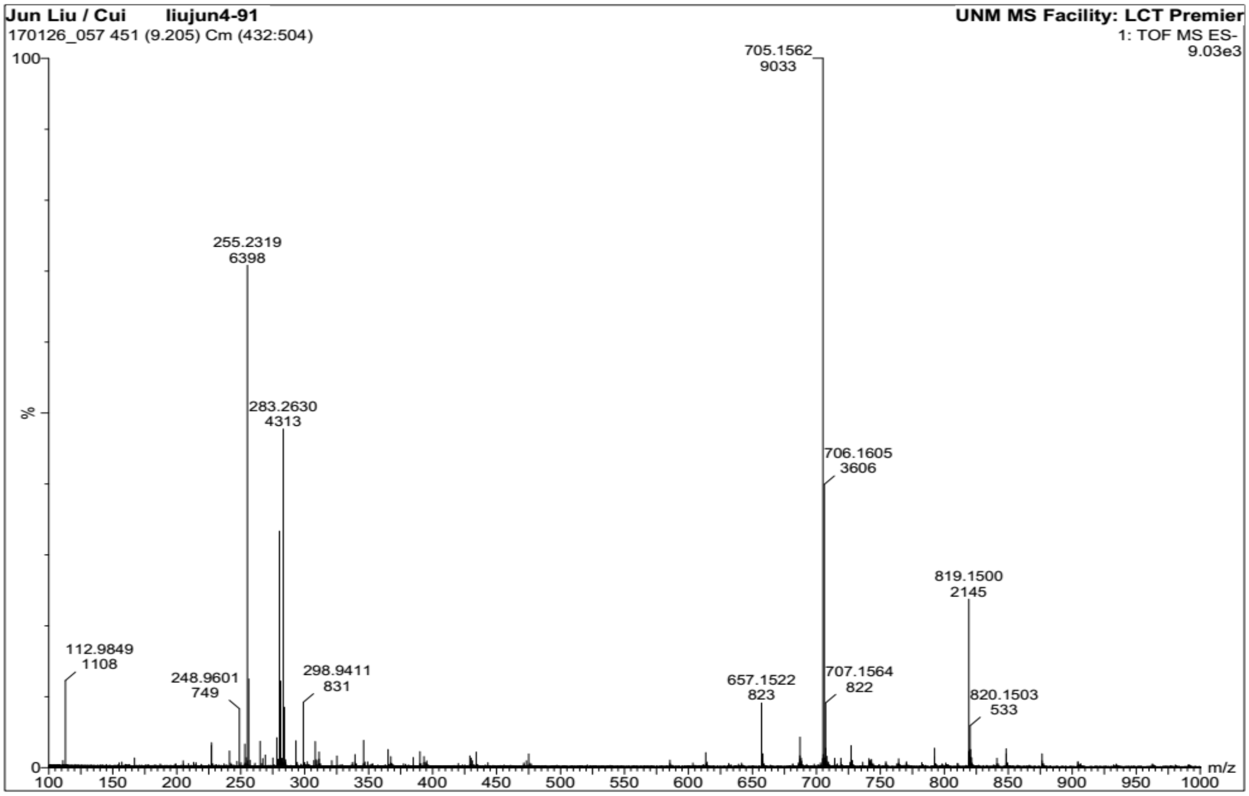
Jun Liu / Cui liujun4-91
170126_057 451 (9.205) Cm (432:504)
UNM MS Facility: LCT Premier
1: TOF MS ES-
9.03e3
705.1562
9033
255.2319
6398
283.2630
4313
706.1605
3606
819.1500
2145
112.9849
1108
248.9601
749
298.9411
831
657.1522
823
707.1564
822
820.1503
533
%
m/z

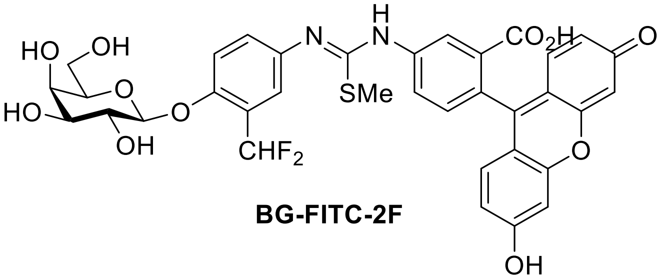

**BG-FITC-2F**

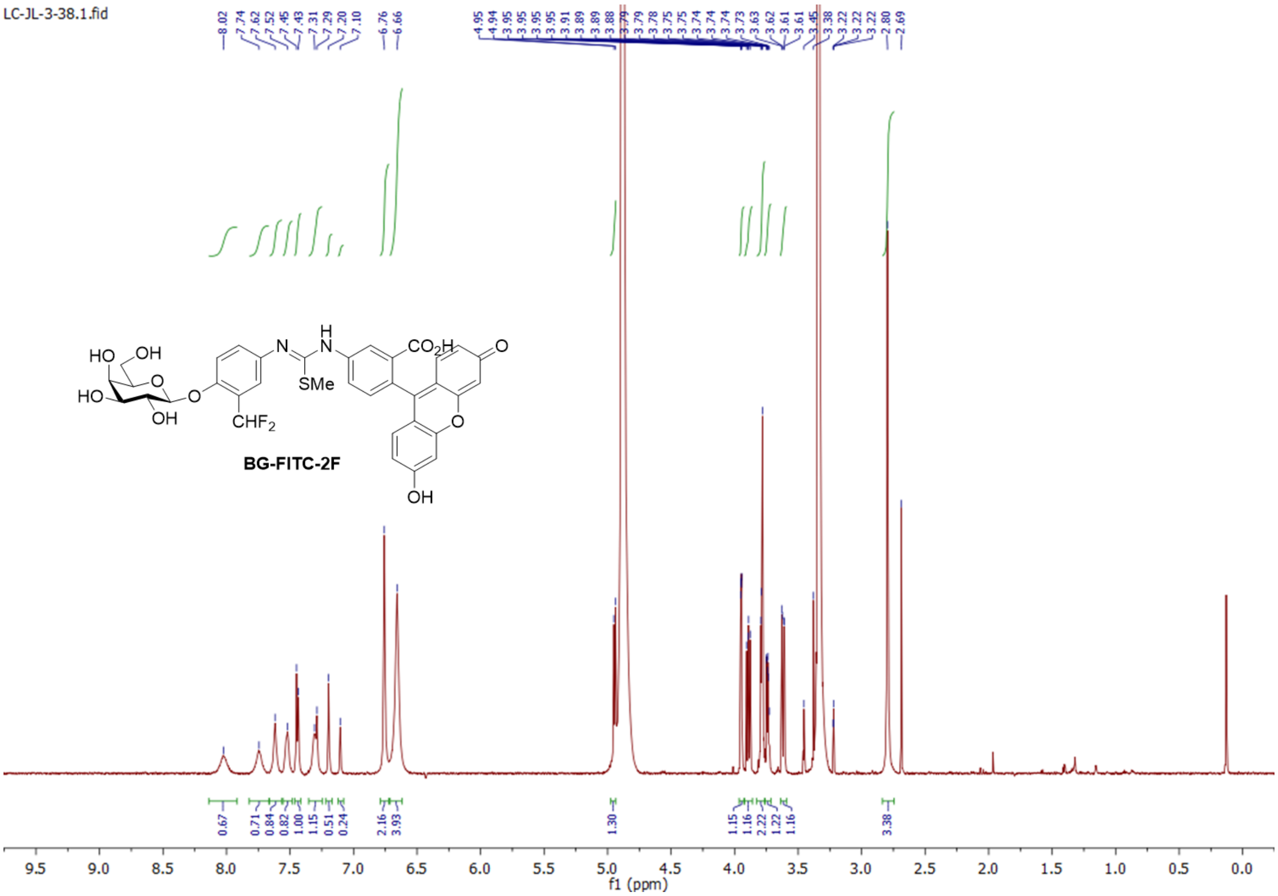

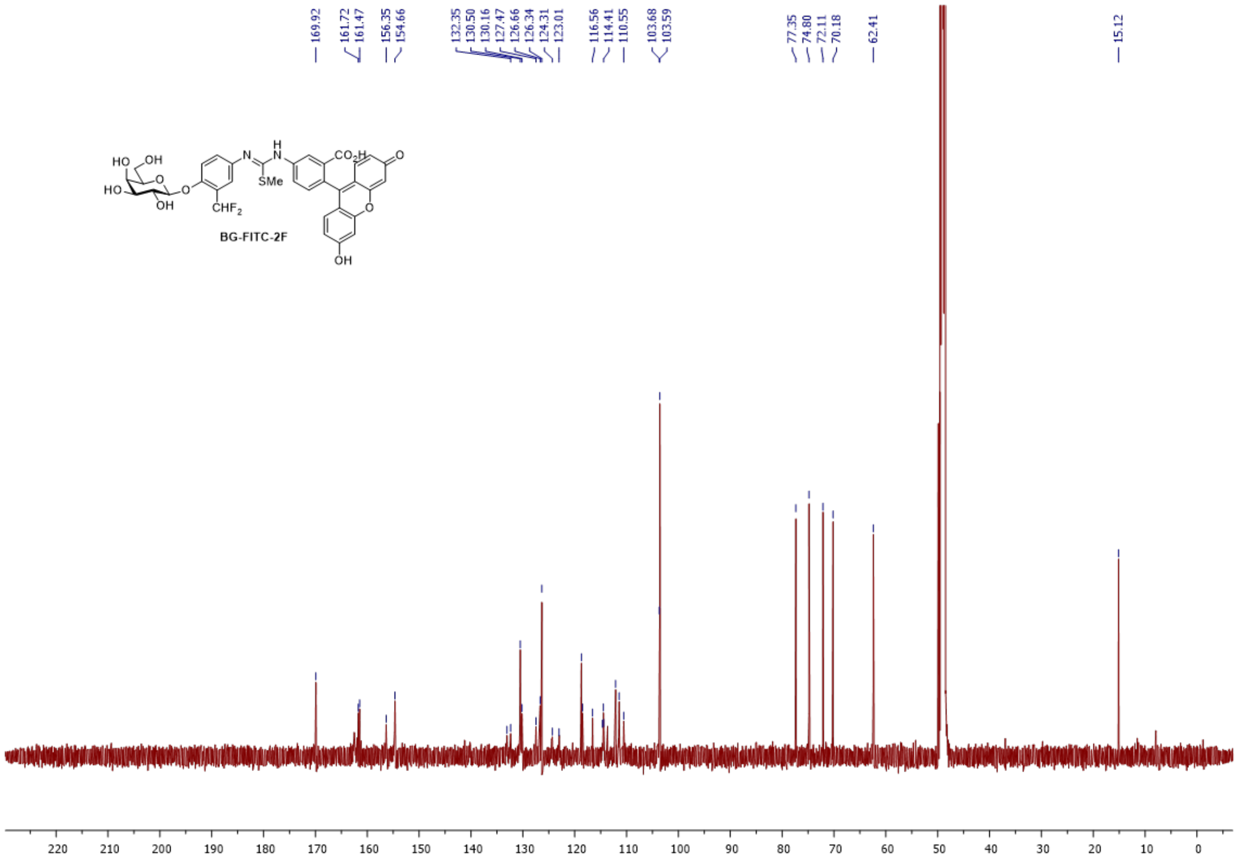
169.92
161.72
161.47
156.35
154.66
132.35
130.50
130.16
127.47
126.66
126.34
124.31
123.01
116.56
114.41
110.55
103.68
103.59
77.35
74.80
72.11
70.18
62.41
15.12
HO
OH
HO
O
OH
O
CHF2
N
H
N
SMe
CO2H
O
O
OH
BG-FITC-2F
220 210 200 190 180 170 160 150 140 130 120 110 100 90 80 70 60 50 40 30 20 10 0

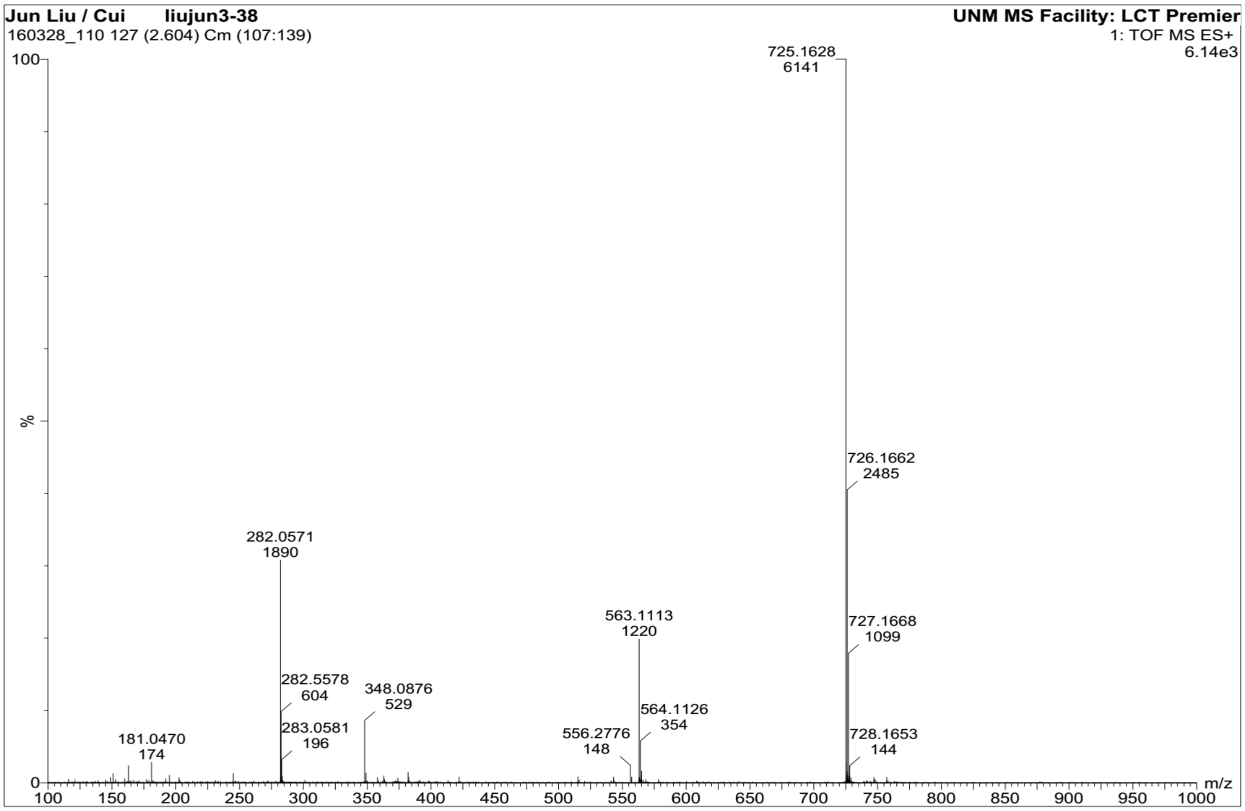
Jun Liu / Cui liujun3-38
160328_110 127 (2.604) Cm (107:139)
UNM MS Facility: LCT Premier
1: TOF MS ES+
6.14e3
725.1628
6141
726.1662
2485
727.1668
1099
728.1653
144
282.0571
1890
282.5578
604
283.0581
196
348.0876
529
563.1113
1220
564.1126
354
556.2776
148
181.0470
174
%
100
0
m/z
100 150 200 250 300 350 400 450 500 550 600 650 700 750 800 850 900 950 1000
